# Experimental Quality Control Induces Changes in Allen Mouse Brain Connectomes

**DOI:** 10.64898/2026.02.20.707091

**Authors:** Vikram Nathan, Stephanie Tullo, Lizette Herrera-Portillo, Gabriel A. Devenyi, Yohan Yee, M. Mallar Chakravarty

## Abstract

The Allen Mouse Brain Connectivity Atlas (AMBCA) is widely used to represent structural connectivity in the mouse brain. The AMBCA consists of tracer injection experiments where neuronal projections axonally connected to the initial injection site are labelled. The resulting whole-brain structural connectomes, derived from a subset of these experiments in C57BL/6 mice, have been used in several studies of connectomic architectures. However, through close inspection of n=437 distinct experiments used in a publicly-available connectome (Knox et al., 2018), we observed experiments with off-target injections, diffuse projections, unrealistically small injections and projections, and anatomical misalignments, affecting the accuracy and applicability of these connectivity experiments. We applied a combined automated and manual quality control (QC) and identified n=56 (∼13% of the original n=437) experiments representing a wide variety of injection and projection failures across the brain. Automated QC was used to detect extreme injection and projection sizes and misalignments, while manual QC was used to detect subtle off-target tracer spreading. Using the remaining n=381 experiments, we rebuilt two different connectomes using previously-published methods; specifically: the regionalized voxel model from Knox et al. (2018), and the homogeneous model from Oh et al. (2014). Our rebuilt connectomes show strong losses in connectivity between regions with limited evidence of structural connectivity by other methods (e.g. hippocampus-medulla, cerebellum-isocortex) and gains in connectivity between regions with strong connectivity evidence (hypothalamus-cerebellum, hypothalamus-isocortex).

Finally, we analyzed the rich club and community organization to demonstrate the potential downstream impacts on the representation of the overall structural connectome architectures of our QC’d connectomes and observed subtle whole-brain organizational changes. We present our rebuilt connectomes, and particularly highlight the regionalized voxel model, as more accurate representations of structural connectivity derived from the AMBCA.

## Introduction

The Allen Mouse Brain Connectivity Atlas (AMBCA) is often used as the gold standard for mouse brain structural connectivity (Oh et al., 2014). In addition to diverse transgenic lines to visualize cell types of interest, the AMBCA consists of serial two-photon microscopy experiments in ∼500 wild-type (WT) adult, male C57BL/6J mice (postnatal days P56 ± 2). Each “experiment” refers to an anatomically distinct injection of a recombinant adeno-associated viral (rAAV) tracer, which expresses enhanced green fluorescent protein (EGFP) under the control of a human synapsin I promoter to specifically label neurons. The tracer spreads anterogradely from the injection site and cannot cross synapses, allowing for the *ex vivo* imaging of the directed, monosynaptic neuronal projections connected to the injection site. The resulting images from these experiments have been used to reconstruct whole-brain axonal connectomes at the regional and voxel levels (Oh et al., 2014; Knox et al., 2018 respectively) that provide significant advantages over commonly used diffusion weighted imaging data acquired using magnetic resonance imaging (MRI): the measured connectivity is direct, rather than probabilistic; all connections are unidirectional and monosynaptic, and the high-resolution of the serial microscopy experiments (0.35 μm) allows for the explicit representation of crossing fibers (Behrens and Sporns, 2012).

These structural connectomes are widely used to interrogate the network architecture of the whole mouse brain. Recent voxel-level anatomical insights include the characterization of hierarchical cortico-thalamic connectivity, an interconnected community recapitulating the default-mode functional network, and differential projections from the thalamic innervations of distinct cerebellar nuclei (Harris et al., 2019; Coletta et al., 2020; Kebschull et al., 2020). These connectomes have also been used to multimodally assess the relationship between monosynaptic connectivity, structural covariance between the thalamus and cortex, and gene expression (Yee et al. 2024; Yee et al., 2018). Because of its anatomical specificity and spatial coverage, the AMBCA has also been used to model whole-brain pathophysiology across a range of pathologies (Mezias et al., 2017; Mezias et al., 2020; Cornblath et al., 2021; Zhou et al., 2025; Torok et al., 2025). A prime example is neurodegenerative disease modelling using the regional connectome from Oh et al. (2014) to simulate the network spreading of pathogenic agents in neurodegenerative diseases, including alpha synucleinopathies and tauopathies, providing mechanistic insights into progressive pathological spreading (Tullo et al., 2025; Rahayel et al., 2022; Henderson et al., 2019; Lubben et al., 2024; Ramirez et al., 2023). In analogous models of pathological spreading in humans, simulations using different connectomic architectures show that modality-specific false positives and negatives can dramatically alter model predictions (Thompson et al., 2024). Therefore, it is critical to have accurate representations of the underlying connectome to understand models of brain organization in the context of brain function and dysfunction.

There are two different connectome models derived from the AMBCA that are commonly used and publicly available: the regional connectome (“homogeneous model”, HM) in Oh et al. (2014), and the voxel-level connectome (“voxel model”) in Knox et al (2018). The HM takes the injection volume from each experiment and assigns the projection to the injected region(s) based on the size of the injection within each region, regardless of the injected location within the region. The voxel model assigns the projection volume to the centroid of the injected region, while inferring connectivity to neighboring voxels using a distance-weighted average. However, while examining the potential use of these connectomes for other projects in our group, we observed experiments with off-target injections, diffuse projections, unrealistically small injections and projections, and anatomical misalignments, in spite of the quality control (QC) procedures documented in their respective publications (Figure 1). These failures may result in the over-or under-representation of connectivity to the injected region.

**Figure 1:**
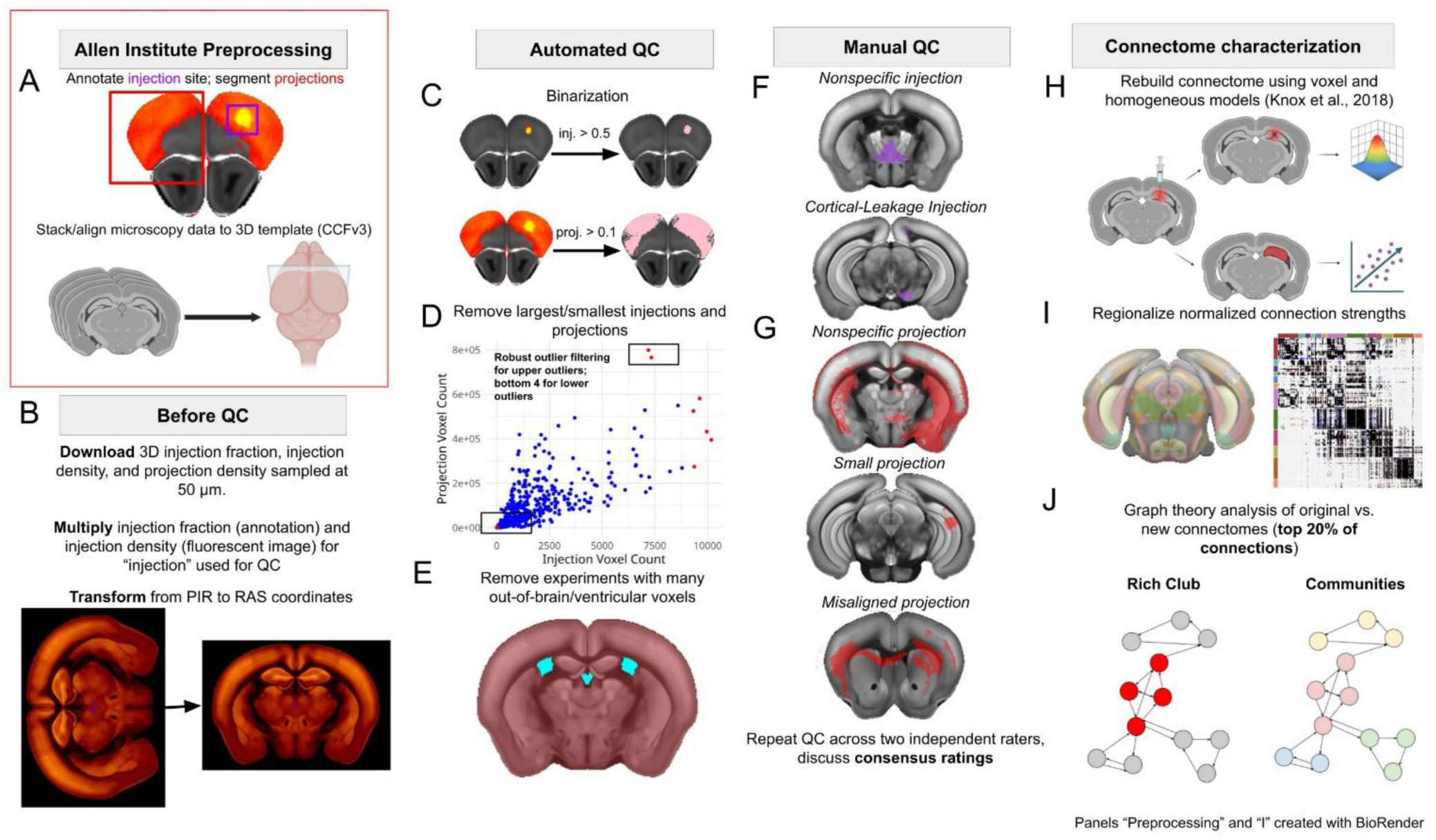
Methods overview. Our workflow consists of three main phases, building off existing 3D-aligned microscopy data from the Allen Institute (A) with additional preprocessing steps (B). 1: Automated QC, including (C) binarization using injection and projection-specific thresholds of 0.5 and 0.1 (D) robust outlier filtering to remove the largest injections and projections, and identifying the smallest four injections and projections (E) removing experiments with the largest and smallest counts of out-of-brain or ventricular voxels, using robust outlier filtering removing experiments with manual QC, and assessment of downstream impacts on the rebuilt structural connectomes. 2: Manual QC for injections (F), including nonspecific injections and cortical-leaking injections; manual QC for projections (G), including nonspecific/widespread, small, and misaligned projection densities. 3: Connectome characterization, including (H) rebuilding the RVM and HM (I) regionalizing the normalized connection strength metric, and (J) rich club and community analyses on the top 20% of connections in the original vs. rebuilt connectomes. For a fully reproducible workflow based on the steps shown above, see https://github.com/vik16nathan/allen_connectome_qc/.

To address this confound, we applied a detailed QC procedure to the n=427 experiments that survived QC in Knox et al. (2018), as well as an additional n=10 WT experiments that were added to the Allen API after 2018. We then rebuilt the homogeneous (HM) and regionalized voxel-model (RVM) connectomes using the n=381 remaining experiments and characterized the impact of using our QC’d connectome on structural connectivity metrics, including the normalized connection strength and top 20% of connections between brain regions. We also examined the downstream impact on graph theory metrics that are commonly used in the connectomics literature, including the rich club and Louvain communities (Van den Heuvel and Sporns, 2011; Gollo et al., 2015; Swanson et al., 2017; Brooks et al, 2024; Coletta et al., 2020; Herrera-Portillo et al., 2025; Ashourvan et al., 2019). We posit that our rebuilt connectomes represent a cleaner, more accurate map of structural connectivity in the mouse brain.

## Methods

### Methods Overview

Our analysis was divided into three main phases: automated QC, manual QC, and characterizing rebuilt whole-brain connectomes after excluding the experiments that did not pass our QC standards (Methods figure). All of our QC’d data was aligned to the Allen Institute’s Common Coordinate Framework, version 3 (CCFv3) template (Wang et al., 2020). We primarily assessed architectural changes in the voxel model, which assigns the connectivity from each experiment to the centroid of the experiment’s injection site, while assuming spatial smoothness for the remaining voxels (Knox et al., 2018). We also characterized the changes in the homogeneous model, which assigns connectivity to parcellated brain regions based on the injection size, regardless of the location of injection within a given brain region (Oh et al., 2014). Finally, we applied graph theoretical metrics to the old vs. new Knox connectome to assess whole-brain organizational changes upon rebuilding the connectome.

### Experimental and 2D Image Processing for Serial Two-Photon Microscopy Data

To measure connectivity to various brain regions, the Allen Institute used iontophoresis to inject a tracer into the cell bodies of various brain regions in wild-type C57BL/6J mice at P56 ± 2 postnatal days. This recombinant adeno-associated viral (rAAV) tracer expresses enhanced green fluorescent protein (EGFP) under the control of a human synapsin I promoter, thus specifically labelling neurons (Harris et al., 2012). The rAAV tracer anterogradely labels axons without crossing synapses; thus, each experiment represents directed, monosynaptic connections to the injection site. The mice were euthanized 21 days following injection before serial two-photon microscopy at a coronal slice interval of 100 μm and within-slice image resolution of 0.35 μm (Oh et al., 2014).

The Allen Institute then processed the scanned two-photon microscopy images. For each experiment, the 2D microscopy images first underwent intensity correction and stitching. The 2D images were then manually quality controlled (QC’d) for specimen and image quality, resulting in the removal of entire experiments with “severe artefacts” (missing tissue, low signal strength, etc.) or the masking of regions with high intensity/frequency artifacts and signal dropout (see Oh et al., 2014). Note that these QC criteria of the 2D images are separate from our QC criteria for the 3D images. After these corrections, the projection pixels in each image were identified by a segmentation algorithm that separated fluorescence from background noise (see Kuan et al., 2015 for more details). After segmentation, the injection site(s) were defined manually in each image by drawing closed polygons around the cell bodies of the infected neurons.

Finally, the Allen Institute stacked these 2D images to form 3D injection and projection images, before being aligned to the Allen Institute’s 3D reference model, the Common Coordinate Framework (Version 3, “CCFv3”), using linear and nonlinear registration.

The aligned 3D injection and projection data (0.35 μm x 0.35 μm x 100 μm) were resampled into isotropic voxels at various resolutions before defining voxel-level metrics that were available to download from the Allen API.

### Defining Injection Density and Projection Density

We downloaded the resampled data at a 50 μm resolution for our analyses, between the finest resolution available (25 μm) and the resolution used to build the connectomes (100 μm). The “injection fraction” was defined as the fraction of 2D pixels in each voxel belonging to the manually annotated injection site, and “injection density” and “projection density” as the sum of segmented injection/projection pixels divided by the sum of all pixels in each voxel. Knox et al. (2018) used the term “injection density” to refer to the product of the injection fraction and injection density values, representing the proportion of segmented neurons in the annotated injection area. We apply the same definition of “injection density” as Knox et al. (2018).

We downloaded the injection fraction, injection density, and projection density from the Allen API for all included (n=427) and excluded (n=63) experiments from Knox et al. (2018). Although Knox et al. (2018) reported n=428 included experiments, experimental data was no longer available to download for tracer 310207648, so this experiment was excluded from all subsequent analyses. We also QC’d ten additional WT experiments that were automatically downloaded by the mouse_connectivity_models package when rebuilding all connectomes (see *Rebuilding the Regionalized Voxel and Homogeneous Model Connectomes*), resulting in a total of n=437 QC’d experiments. The n=63 experiments removed in Knox et al. were manually excluded for a variety of reasons, including (1) injections located in multiple major divisions (see Supplementary Table 1 for list of major divisions) (2) subcortical injections with cortical leakage (3) extremely small projections. We will extend these criteria to more stringently QC the experiments that passed Knox et al.’s evaluation.

### Transforming and Thresholding Injection Density and Projection Density

After downloading the injection and projection data, we converted data from the PIR (+x: Posterior-Anterior, +y: Inferior-Superior, +z: Right-Left) space used by the Allen institute to RAS (+x: Right-Left, +y: Anterior-Posterior, +z: Superior-Inferior) coordinates. RAS coordinates are the canonical orientation in preclinical imaging, under which positive directions correspond to right (x), anterior (y), and superior (z); this coordinate system is also compatible with our existing image analysis pipelines (Desrosiers-Grégoire et al., 2024; Germann et al., 2025). We rotated the axes and redefined the position of the origin to the center of the anterior commissure as it crosses the sagittal midplane, consistent with the standard DSURQE MRI atlas (Dorr et al., 2008; Steadman et al., 2014; Ullmann et al., 2013; Richards et al., 2011; Qiu et al., 2018). For anatomical analyses of the injection and projection data, we downloaded the most recent CCFv3 template (Wang et al., 2020) and label file from the Allen API and repeated the above transformation. Additionally, we created a brain mask from the nonzero values in the label file. We apply the exact same mathematical conversion, which consists of rotations and dilations, to all CCF templates, injection and projection data (https://github.com/vik16nathan/allen_connectome_qc/blob/main/before_manual_qc/transform_space.py); hence, all relationships between the template and experimental data are maintained following the PIR-RAS conversion.

We sought to ensure that any QC failures were not simply due to the lowest injection and projection values, which could represent poorly segmented projection data or autofluorescence. To this end, our automated and manual QC only considered injection density values over 0.5 and projection density values over 0.1 (Figure 1A), values that were heuristically chosen after visual inspection to retain the majority of the injection and projection data while eliminating noise. These thresholds were similar to the threshold of 0.2 applied to the projection density data within the *Brain Explorer* application used to visualize the Allen Brain Connectivity Atlas (Feng et al., 2015). We note that the projection QC issues identified in this analysis can also be seen in the Allen Brain Explorer (see https://connectivity.brain-map.org/). These binarized files were used for automated QC, and to generate slice images for manual QC. Note that when rebuilding the connectome, we used the full, unthresholded data (see *Rebuilding the Regionalized Voxel and Homogeneous Models*).

We did not interpolate or resample any of the downloaded injection or projection data, and did not apply any additional masking or preprocessing steps beyond the PIR → RAS conversion and binarization. Since thresholding was the only transformation applied to these data before building the original connectomes, our QC evaluated the data in the format used for downstream connectivity modelling (Knox et al., 2018).

### Automated Quality Control

Before visually inspecting all experiments, we defined automated exclusion criteria. To this end, our automated and manual QC only considered injection density values over 0.5 (as was done in Knox et al.) and projection density values over 0.1 (Figure 1A). Both of these values were heuristically chosen after visual inspection to retain the majority of the injection and projection data while eliminating noise. We first excluded experiments with the largest numbers of out-of-brain and ventricular projection voxels, using the CCFv3 brain mask and ventricular region labels in the CCFv3 label file to identify failed experiments (Figure 1B). These failed experiments represent alignment failures when stacking or registering the 2D microscopy data to the 3D template, particularly evident with corpus callosum misalignment, or QC failures with the tracer spreading nonspecifically throughout the brain (Figure 1F). For our ventricle analysis, we did not consider voxels in the third ventricle and cerebral aqueduct, as these regions were too thin to robustly capture misaligned experiments. We then examined the distributions of the number of out-of-brain and ventricular projection voxels, identifying outliers outside the following interval, using the formula below from Hubert and Vandervieren, 2008:

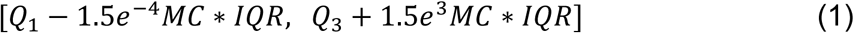

Where Q_1_ and Q_3_ refer to the first and third quartile, and MC refers to the medcouple, defined for univariate sample {x_1_,…, x_n_} from a continuous unimodal distribution F:

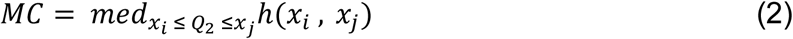

With Q_2_ as the sample median, and the kernel function *h* defined for all x_i_ ≠ x_j_ as:

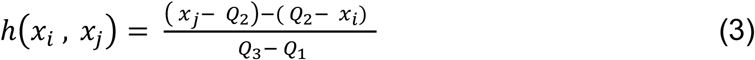

This formula applies exponential right skewness adjustment coefficients multiplied by the values 1.5 times the interquartile range (IQR) from the median.

For the full implementation of this outlier filtering, using the robustbase package (Maechler et al., 2025), see https://github.com/vik16nathan/allen_connectome_qc/blob/main/after_manual_qc/overall_qc_exclusion.R.

We note that this formula identifies no outliers for ventricular voxel counts and only upper outliers for out-of-brain voxel counts, using 50 μm isotropic voxels (threshold for exclusion: ≥ 10,230 out-of-brain voxels).

Secondly, after excluding experiments with out-of-brain and ventricular voxels, we then excluded experiments with the largest and smallest injection and projection voxel counts (Figure 1C). Although it is possible for experiments to have very large or small injections and projections, we argue that the downstream assignment of these connectivity patterns to individual brain regions or voxels is biologically implausible, resulting in false-positive or false-negative connections. We used the same right-skewness-adjusted robust outlier filtering criteria, which only identified upper outliers for injection and projection voxel counts (injection exclusion threshold: ≥ 9,328 voxels; projection exclusion threshold: ≥ 581,630 voxels). We also excluded experiments with the bottom four smallest injection and projection voxel counts (injection: ≤ 4 voxels; projection ≤ 343 voxels).

### Manual Quality Control

We then manually QC’d all n=437 experiments’ injection and projection densities to detect nuanced QC failures. For example, an implausibly small projection from a well-connected brain region like the hippocampus would still be larger than a biologically realistic cerebellar projection; hence, “small” vs. “nonspecific” injection and projection volumes depend on the region of injection. To this end, we QC’d all images using PyQC (see https://github.com/CoBrALab/ for make_slice_images and PyQC), evaluating the injection and projection data from each experiment atop the CCFv3 template using a consistent set of slice series. Applying the default parameters in make_slice_images.sh and no cropping, we generated ten equidistant x, y, and z slice images for each injection and projection volume, resulting in a total of 30 images per volume. For injection or projection volumes with no voxels on the chosen slices, we generated cropped slice images based on the location of the injection or projection. For full QC images, see Supplementary Data 1 (https://zenodo.org/records/20276701). Visual inspection of a single experiment’s data atop brain slices with PyQC allowed us to qualify whether the size, spatial distribution, and alignment of a given injection or projection was sensible given the underlying anatomical structures, based on manual QC criteria for injections and projections outlined in the following paragraph (see Supplementary Methods Figure 1 for examples).

We then applied separate QC criteria to the injection and projection images. For all images, we defined a rating of “0” as a “pass,” and a rating of “4” as a partial failure not severe enough for removal. For the injection images, a rating of “1” (Figure 1D) denotes a large or nonspecific injection spanning multiple brain regions. A rating of “2” (Figure 1E) denotes subcortical injections with substantial cortical leakage, or, conversely, a cortical injection that also infects subcortical areas, generalizing the criteria from Knox et al. (2018). For the projection images, “1” (Figure 1F) denotes nonspecific tracer spreading to off-target regions spanning multiple major brain divisions, “2” (Figure 1G) denotes little or no long-distance projections from the injection site (outlined in Knox et al.), and “3” (Figure 1H) denotes a misalignment to the anatomical template. We removed all experiments that met these criteria before rebuilding the RVM and HM connectomes (see Supplementary Data 2 for full list of all numerical ratings).

Note that we also defined a rating of “5” for projection data from an experiment with a very small number of out-of-brain voxels that is not necessarily misaligned, and a rating of “9” for an experiment with no injection or projection data in the original ten-slice QC image, prompting further investigation and the generation of a cropped image. The “9” images did not necessarily have the smallest number of projection voxels, but often had voxels between the default slices generated by PyQC. We also repeated our manual QC on all injection and projection data with lower binarization thresholds (0.01) (see Supplementary Data 1), and observed ratings of “5” in experiments with voxels that aligned well with the anatomy of the underlying template, indicating possible partial-volume effects that could be masked out, rather than a registration failure. Therefore, we did not define “5” experiments (n=1) as QC failures.

Two independent raters (V.N, S.T.) evaluated all injection images at a 0.5 binarization threshold and projection images at a 0.1 threshold before meeting to discuss consensus ratings on any disagreements. The final consensus QC ratings for all injection and projection images were used to define which experiments to exclude from further analysis. Additionally, V.N. separately examined manual QC images at a 0.01 threshold to confirm that the designation of “failed” experiments was not threshold-dependent. For the initial disagreements and full ratings for all experiments, see Supplementary Data 2.

### Rebuilding the Regionalized Voxel and Homogeneous Model Connectomes

After removing experiments that did not pass our QC criteria, we characterized the impact of these removals on canonical models of whole-brain connectivity. The main connectome evaluated was the voxel-level connectome from Knox et al. (2018). This model estimates connectivity by assigning each experiment’s projection strengths to the centroid of the injection site, and infers connectivity to voxels neighbouring the injection site using a Gaussian kernel average across experiments within a single major brain division (see Supplementary Table 1 for list of all brain divisions). Using code adapted from Knox et al., 2018, we then rebuilt the voxel-level model and calculated a dot product to “regionalize” the connection strengths over n=291 gray matter regions, resulting in the regionalized voxel model (RVM).

We also rebuilt the HM connectome from Knox et al. (2018), which builds a regional connectome using a separate set of mathematical assumptions defined in the “homogeneous model” used by Oh et al. (2014). Algorithmically, the HM uses nonnegative linear regression to calculate the “weight” of connections between a source region and a target region, by predicting the projections in a target region from the injections in a source region. The HM relies on two key mathematical assumptions: “additivity,” which assigns each experiment’s projection strengths to all regions circumscribing the experiment’s injection site, and “homogeneity,” which learns a single connectivity weight from a source to a target region, regardless of the location of the corresponding injections (“homogeneity”). However, the original HM is computed for n=291 regions, whereas the connectome from Oh et al. (2014) is computed on a broader set of n=213 regions. Thus, we modified Knox et al.’s code for the HM reconstruction to only model connectivity to n=211 mesoscale regions, specifically excluding the dorsal and ventral subiculum that were no longer available within the most recent version of the CCFv3 atlas (Wang et al., 2020).

To rebuild the RVM and HM connectomes, we reran the algorithms located within the Allen Institute’s *mouse_connectivity_models* package (https://github.com/AllenInstitute/mouse_connectivity_models), which accessed all n=498 WT experiments in the current version of the Allen API (accessed February 2026) within the *allensdk* package (version 2.16.2, see https://github.com/AllenInstitute/AllenSDK/releases), but excluded the n=63 experiments originally excluded by Knox et al. (2018). This resulted in a total of n=435 experiments used to rebuild all connectomes before our QC. It is important to note that although we QC’d experiments 100142354 and 112670853 due to their inclusion in the Knox connectome, they were not used for the original or rebuilt RVM or HM connectomes described in this paper.

We rebuilt the RVM and HM connectomes using the n=381 remaining experiments. We kept all connectivity modelling algorithms the same as in Knox et al., but retrained the HM weights and RVM kernel regression smoothness parameters (*σ)* to reflect our newly QC’d list of experiments. For the HM (build_homogeneous_model.py), which involves nonnegative linear regression, the only parameter that we changed was the list of experiments excluded. However, for the RVM, we first retrained the smoothness parameters for the fitted Gaussian kernel used to estimate connectivity within each major division (run_hyperparameter_selection.py). We then applied these parameters to the QC’ed set of experiments to estimate connectivity between brain regions (build_model.py). By default, these algorithms produce maps of “normalized connection strengths,” which represent the raw connection strength between any two regions divided by the size of each region.

### Nested Leave-One-Out Cross-Validation Analysis

To assess the sensitivity of the retrained connectivity models, we replicated the leave-one-out cross validation analyses using code from Knox et al., but repeated on the *m* experiments within each major division that survived QC (https://github.com/AllenInstitute/mouse_connectivity_models/tree/master/paper/figures/model_comparison; see run_nested_homogeneous.py, run_nested_voxel.py, and compile_table.py). The error metric used is MSE_rel_, which is the mean-squared error divided by the average squared norm of the prediction and left-out data.

First, a single experiment is held out in the outer loop. Then, in the inner loop, *m* − 1 different models are fitted on *m* − 2 experiments to predict a second held-out-experiment; the training error is calculated for each of these models. In the case of the voxel model and RVM, these *m - 1* models are compared to identify the optimal hyperparameter *σ* of the Gaussian kernel function; for the HM, the optimally-predictive nonnegative regression weights are identified. Finally, the identified “best model” from the inner loop is used to predict the connectivity to the held-out experiment in the outer loop; this is the testing error. This process is repeated across all *m* experiments in each division, and the errors are averaged.

This cross-validation procedure is also repeated on the subset of the *m* experiments that have at least one other experiment with the same injected region, which Knox et al. defines as the experiments with the “power to predict” (PTP).”

### Rich Club and Community Detection Analysis

Finally, we provided examples of how our QC impacts graph theoretical metrics of network-level organization. We examined the top 20% of overall connections across the RVM. Whichever connections remained were then assigned as “edges” to the corresponding source and target regions, which were the “nodes” in our network. We chose two metrics: the topological rich club nodes (based on the continuous rich club coefficient for each node) and the Louvain community assignments for each node.

These metrics complementarily capture which nodes are well-connected among themselves and to the rest of the network (rich club), and which groupings of nodes are well-connected between themselves (communities) (see schematic in Figure 1).

Our rich club analysis considered both the continuous, nodal “rich club percentage,” or the proportion of all rich clubs that a given brain region is involved in, as well as the global “topological rich club” defining the most highly interconnected regions (see Supplementary Methods). Our community analysis assigned a single, discrete community value to each node, based on the most stable consensus assignment of communities across resolution parameters affecting the size of the data-driven communities. For all analyses, we used the Brain Connectivity Toolbox (version 2019-03-03) in MATLAB (version R2024b) (Rubinov et al., 2009). Our approach was inspired by similar work that characterized these metrics using the Knox et al. connectome (Herrera-Portillo et al., 2025; Coletta et al., 2020); for algorithmic details, see *Rich Club Algorithm* and *Community Detection Algorithm* in Supplementary Methods.

## Results

### Automated QC Reveals Registration and QC failures

We identified several experiments as outliers for the number of projection voxels outside the brain (n=3), but no experiments as outliers for ventricular voxels. We note that one of these outliers, experiment 112670863, was included in the original RVM connectome but was no longer available through the *allensdk* package. Further inspection of the projection patterns of the top ‘out-of-brain’ experiments revealed misalignments to the underlying anatomical template and key visual landmarks like the corpus callosum (Figure 2A). Despite not being identified as outliers after skewness adjustments, the top two experiments for ventricular voxels were identified as outliers for the largest overall projection volumes, indicating that the number of ventricular voxels could capture diffuse, non-specific tracer spreading (Figure 2A, 2B).

**Figure 2:**
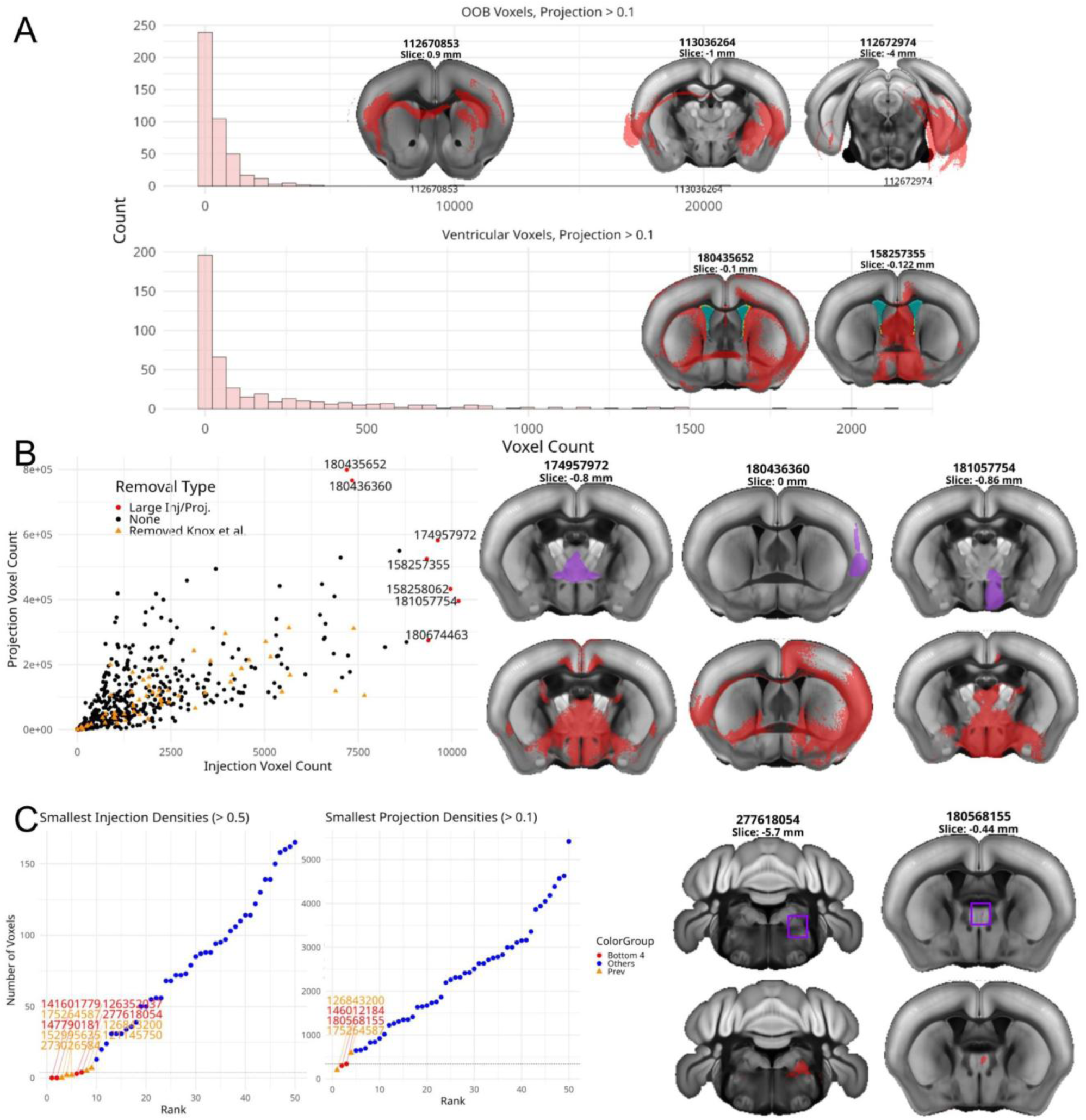
Automated QC results identify many QC failures, including: (A) experiments that are significant outliers for out-of-brain voxel counts (after robust outlier filtering), and severe experiments for the number of ventricular voxels, indicating misalignment and/or nonspecific tracer spreading (B) experiments that contain the highest number of injection and projection voxel points, overrepresenting connectivity to injected regions and biasing the connectome with nonspecific tracer spreading and (C) experiments with the lowest number of injection and projection voxel counts, which inflate connectivity derived from sparse neuronal subpopulations that are likely unrepresentative of regional or voxel-level connectivity. Note that all counts are calculated on the thresholded, binarized data.

We also identified upper and lower extrema for injection and projection voxel counts (Fig. 2B and 2C, respectively) and compared the binarized injection and projection voxel counts to experiments that failed the QC criteria in Knox et al. (2018). Importantly, we see that our identified outliers for the largest projections, largest injections, and smallest injections are more extreme than those identified in Knox et al. (2018), and that our excluded experiments for smallest projections are within 100 voxels of those excluded by Knox et al. (2018) (Fig. 2C).

In total, after these automated QC steps, we removed n=17 experiments (Supplementary Figure 1). However, we also observed that the n=63 experiments removed by a separate manual QC at the Allen Institute show a diverse range of injection and projection voxel counts, indicating more nuanced QC failures that were specific to the injected region (Fig. 2B). Thus, our own manual QC on the n=435 analyzed experiments was necessary to characterize the remaining failure modes (Figure 1).

### Manual QC Reveals Spatially and Qualitatively Diverse Failures

Two independent raters applied our manual injection and projection QC criteria to all n=437 experiments prior to deciding on consensus ratings (see *Methods*). We observe a high inter-rater reliability in the injection QC criteria (86%) and projection QC criteria (86%) before consensus adjustment. Despite the high percent agreement, we see only moderate values of Cohen’s kappa (injection = 0.288; projection = 0.174), since S.T. was stricter than V.N. when applying the criteria (see Supplementary Table 4 for initial ratings distribution). Hence, harmonized ratings following discussion between the two raters were necessary to consistently identify failures.

Our consensus ratings resulted in the manual removal of n=50 experiments. Taken altogether with our automated QC, we remove n=6 experiments based on Automated QC alone, n=39 experiments with manual QC alone, and n=11 experiments based on both manual and automated QC. We note that two experiments from Knox et al. were already excluded when downloading experiments from the updated Allen API; hence, our QC excluded n=54 novel experiments from the n=435 downloaded experiments, resulting in n=381 experiments used to rebuild all connectomes.

We observe a wide variety of experiments removed across major brain divisions and failure modalities (Figure 3C), with the greatest number of removed experiments with injections in the hypothalamus, hippocampus, and midbrain. We see that manual QC distinctly captures cortical leaking injections, nonspecific injections, and small projections from the injected region (Supplementary Figure 1). On the other hand, we see that automated QC is the most effective at identifying nonspecific and misaligned projections. Many of these most severe experiments fail both automated and manual QC criteria, providing compelling evidence for removal of these experiments and the validity of our criteria. We also noticed consistencies in failure modes across major brain divisions. For example, all failed cerebellar experiments had small projections, and all but two failed hypothalamic and midbrain experiments had off-target injections.

**Figure 3:**
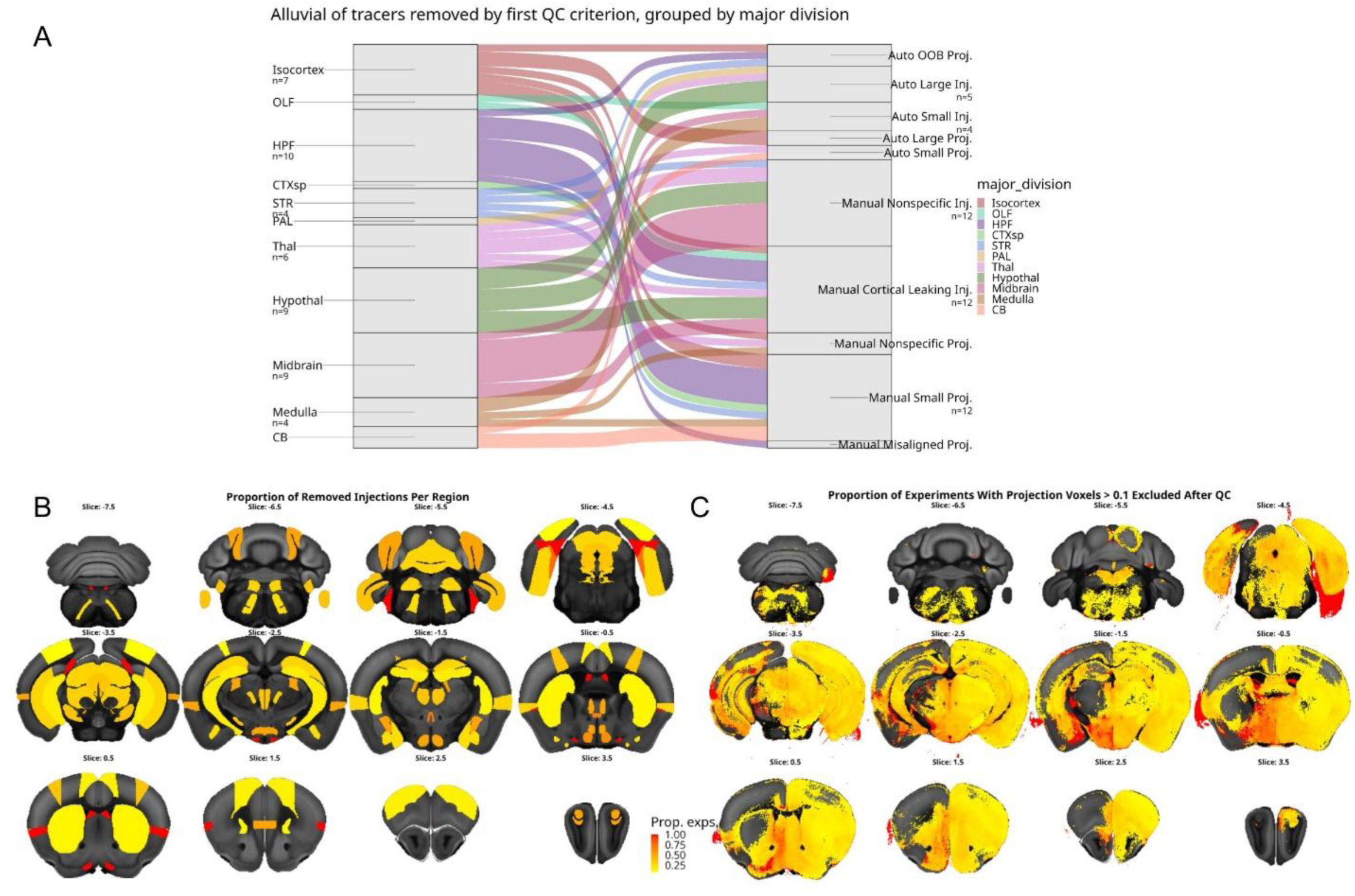
We show the full set of manual/automated QC Removals. (A) All excluded experiments are organized by the first exclusion criteria (see Supplementary Figure 1) and the major division injected (abbreviations in Supplementary Table 1), with the number of experiments labelled for categories with greater than three experiments. (B) The proportion of removed injections in each region of the CCFv3 atlas, relative to the total number of experiments that injected each region. (C) The proportion of removed projections in each voxel, relative to the total number of projections in each voxel. The total number of injections/projections in each region/voxel are calculated over all n=437 experiments.

Nonetheless, for the hippocampal formation, we also observed a mix of connectivity failures, including out-of-brain projections, cortical leaking injections, small projections (n=5), and misaligned projections. For a more detailed analysis of these experiments, see *Extended Description: Manual QC Failures* within the Supplementary Information.

We then examined the locations of the lost connectivity. We find that we lose all injections into small cerebellar subregions and brainstem nuclei, which could potentially result in false-negative connectivity to these regions; however, since all other regions have at least one remaining injection, our procedure also has the potential to reduce false-positive connections (Fig. 3B) As expected, we lose 100% of projection voxels outside of the brain and within the medial ventricle, but also lose large proportions (∼75%) of projections to hypothalamic areas (Fig. 3C). Still, we see a loss in projection voxels across the brain, indicating that our QC removals are not specific to a single brain region.

### Continuous Architectural Changes in Rebuilt Connectomes

After removing the n=56 experiments, we rebuilt the connectivity strengths between n=291 regions in the RVM and n=211 regions in the HM connectome (see *Methods*) (Knox et al. 2018; Oh et al., 2014). We observed minimal changes in the distribution of logged, maximum-normalized connectivity strengths in the RVM and the HM (Supplementary Figure 3). However, we also observed that our QC results in meaningful reorganizations of connectivity for both the RVM and HM connectomes (Supplementary Figures 3C and 3D). Most strikingly, due the nonnegative least-squares regression algorithm used for the HM connectome, we see ∼20,000 regional connections, out of 89,042 connections, that are assigned a strength of “0” before QC but nonzero connections after QC, and vice versa for an additional ∼20,000 connections. On the other hand, for the RVM connectome, we see a stronger agreement between the pre-QC and post-QC connection strengths (Spearman ρ=0.978), albeit with meaningful reorganizations in connectivity, represented by strong deviations from the line of unity in the rank correlation plot. In particular, we see a large number of connections with strongly reduced ranks post-QC versus pre-QC (far below the line of unity), indicating potential losses in false-positive connectivity.

We then z-scored the “normalized connection strength” in the original vs. rebuilt connectomes, before calculating the difference in z-scored connectivity in the new minus the old RVM and HM connectomes. This procedure reveals strong gains and losses in the z-scored connection strengths between regions in the RVM and HM connectomes following QC (Figure 4A and Supplemental Figure 4A, respectively). Consistent with the distributional changes, we see that the HM shows stronger losses (negative differences) in connectivity, whereas the RVM connectome is more robust to the removed experiments; hence, our main results will focus on the RVM connectome. In the RVM connectome, we observed that the largest losses in z-scored connectivity strength were in hypothalamus-hypothalamus connections (Figure 4A, #1), particularly contralaterally. We also observed strong losses in connection strength between medullary regions and contralateral thalamic (#2) and midbrain (#3) regions, and between isocortical regions and the contralateral olfactory bulb (#4), corticular subplate (#5), and striatum (#6). However, especially for hypothalamus-hypothalamus connections, these strong losses in connectivity strength were often accompanied by a strong gain in connectivity to nearby regions (#7). We interpret the losses in connectivity as the removal of false-positive connections, and the gains in connectivity as a redistribution of modeled connection strengths after the removal of “noisy” experiments with false-positive and false-negative connectivity patterns.

**Figure 4:**
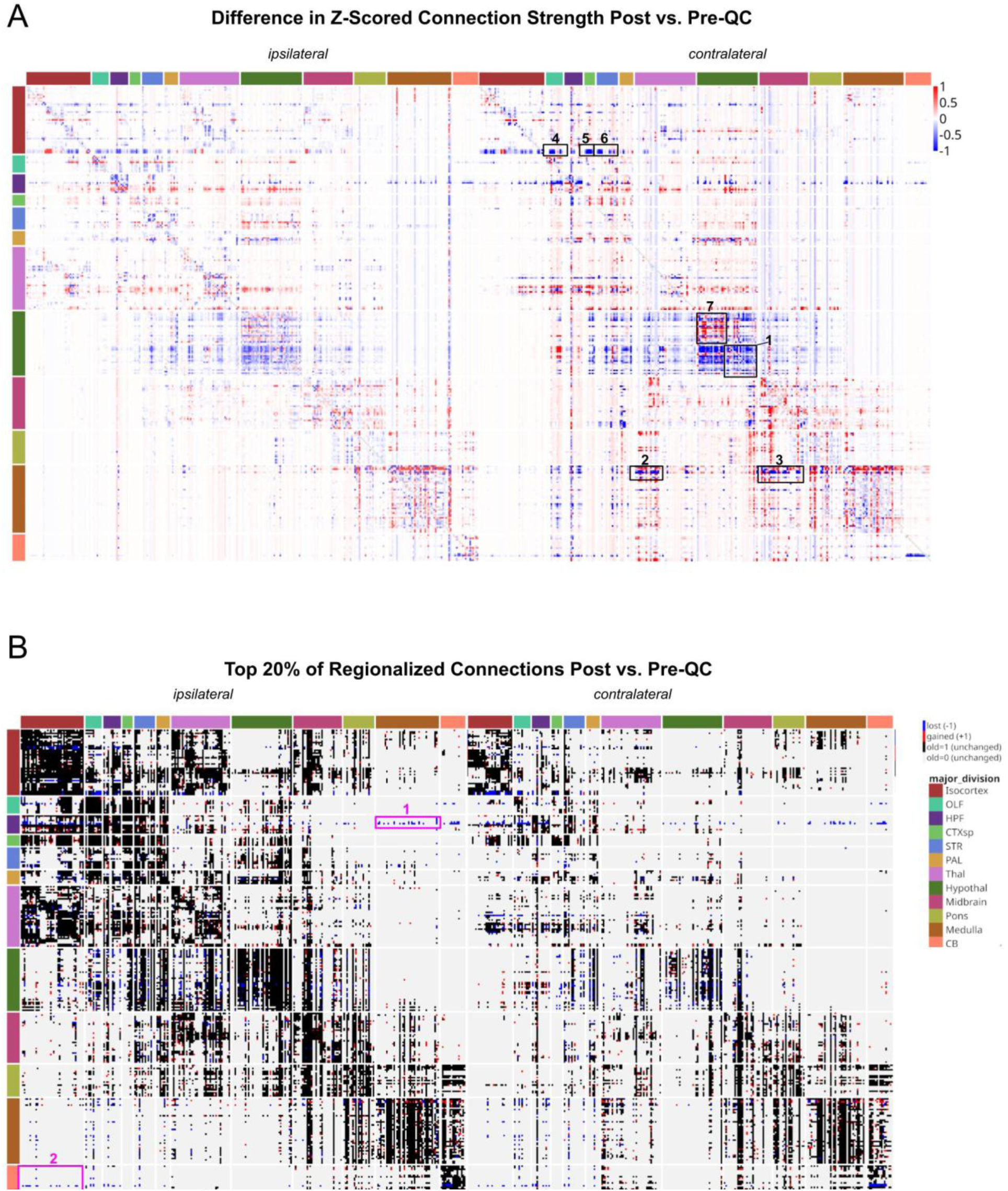
We observe whole-brain architectural changes in the RVM (n=291 regions) after QC, when examining the (A) difference in z-scored, normalized connection strengths, thresholded from [-1,1] (B) discretely, when examining the lost (blue), gained (red) and unchanged (black/gray) connections within the subset of the top 20% of the old vs. rebuilt connectomes. The numerical labels correspond to specific gains or losses in connectivity mentioned in the *Results*.

We thus examined the overall change in regionalized connection percentile aggregated across major brain divisions. We examined the percentile rather than the z-score due to the wide use of percentile-based thresholding to enforce a network with a given density for connectomic analyses (Fornito et al., 2013). In the RVM connectome, we see the strongest losses in the connectivity of the hypothalamus to the contralateral cortical subplate (-9.41%) and the hippocampus to ipsilateral medulla (-8.25%) (Supplementary Figure 5A). Additionally, we observed widespread losses in cerebellum-forebrain connectivity (isocortex, olfactory bulb, hippocampal formation, corticular subplate), but a strong gain in connectivity to hindbrain divisions (pons, medulla). Ipsilaterally, we also observed a general loss of connectivity to the hypothalamus, especially to the cortical subplate (-3.91%). These changes were consistent with the changes in the HM (Supplementary Figure 6A), especially the strong losses in hippocampus-cerebellum (-4.67%) and hypothalamus-cortical subplate connectivity (-1.64%). Nonetheless, each model shows distinct gains in connectivity; namely ipsilateral cerebellum-pons in the voxel model (+2.45%) and ipsilateral midbrain-cortical subplate in the HM (+2.69%).

### Discrete Architectural Changes in Rebuilt, Thresholded Connectomes

After characterizing continuous connectivity changes across all connections, we wanted to assess the impact of our QC on the strongest connections, as these are more likely to represent true-positive monosynaptic connections between brain regions. We binarized our connectomes before and after QC using two separate thresholds that retained the top 20% and top 5% of connection strengths, inspired by the density of remaining connections after applying high versus low p-value thresholds, as determined by Oh et al. (2014) (Zalesky et al., 2016). We then visualized the changes in the binarized region-region connectomes.

Although the majority of the strongest connections in the RVM and HM connectomes were retained, at a 20% threshold, we observed brain regions with large numbers of lost and gained binary connections within each model (Figure 4B and Supplementary Figure 4B, respectively). Additionally, we observed more subtle losses and gains in connectivity when comparing the top 5% of connections across both models pre and post-QC (Supplementary Figure 7). Most strikingly, in the RVM connectome, we observed a large number of lost connections between a hippocampal region (identified as the *induseum griseum*) and between the medulla and cerebellum (Figure 4B, #1), as well as between the cerebellum and isocortex (#2). We observed similar losses in hippocampal connectivity within the top 5% of connections in the RVM (Supplementary Figure 7A).

When aggregating the top 20% of connections across major divisions, after QC, 100% of previous connections between the hippocampus and both the ipsilateral and contralateral medulla were lost (Supplementary Figure 5B). Similarly, all bilateral connections from the olfactory bulb to the cerebellum were lost, and most of the connections between the cerebellum and isocortex were lost (94% ipsi; 100% contra). In the HM, we also observed a 100% loss in hippocampus-medulla and 81% loss in cerebellum-isocortex connections (Supplementary Figure 6B). We also observed a complete loss in the top 5% of connections in the RVM between the hippocampus and medulla (Supplementary Figure 8A).

Our discrete gains in connectivity, after percentile-based thresholding, are based on a reordering of overall connection strengths after removing false-positive connections and refitting the connectivity models. For example, in the midbrain, after removing a number of experiments with nonspecific injections, in the RVM, we observe a 700% increase the top 20% of connections between the midbrain and ipsilateral cerebellum, a trend that is mirrored by a 55% gain and 5% loss in existing connections in the HM (Supplementary Figure 5B, 6B). Similarly, when examining the larger set of connections between the hypothalamus and contralateral isocortex, we observed a 95% gain in the RVM and a 178% gain/6% loss in original connections in the HM. We also observed general gains in hypothalamus-contralateral cerebellum connectivity (300% gain RVM; 100% gain and 22% loss in HM). Nonetheless, these trends were not present in the top 5% of connections in the HM after QC; we instead observed a general increase in connectivity from the hypothalamus to all major divisions except the cerebellum, a trend that is not present in the RVM (Supplementary Figure 7; Supplementary Figure 8).

Taken together with the continuous changes shown in the previous section, we claim that our QC criteria removes false-positive connections while preserving and adding biologically meaningful true-positive connectivity patterns, a trend that is particularly evident when examining the binarized connectomes from the RVM. However, we acknowledge that our QC is dependent on multiple heuristic parameter choices.

Therefore, we run a sensitivity analysis on (1) injection and projection volume binarization thresholds for automated QC, (2) cutoffs for the number of lower outliers (3) automated-only vs. full QC (4) the binary “losses” in connectivity across the top 15, 20, and 25% of connections (see *Sensitivity Analysis of QC Across Parameters* in Supplementary Information, and Supplementary Figure 10). We show that the downstream reconstructed connections in the RVM and HM are robust to slight QC parameter changes, with the RVM remaining particularly robust (see *Discussion*).

### QC Reduces Leave-One-Out Voxel Model Error in Six Major Divisions

To assess the ability of the QC’d voxel model, RVM, and HM to predict held-out connectivity, we ran a nested leave-one-out cross-validation analysis using our n=381 experiments after QC. Comparing our results against the results from Knox et al., 2018, we note lower mean-squared errors (MSEs) across some, but not all major divisions (Supplementary Table 3). Nonetheless, we observe lower MSEs in the isocortex, olfactory, hippocampal, cortical subplate, medulla, and cerebellar regions within the voxel model (before regionalization). Our reduction in error for the isocortex is particularly encouraging given that the isocortex contained the largest number of experiments before and after QC (n=128; n=121, Supplementary Table 2), indicating that kernel regression is more capable of predicting the voxel-level connectivity patterns to the remaining n=121 experiments than the original n=128. Additionally, for the isocortex, olfactory areas, hippocampus, and medulla, we also see reductions in the MSE for the regionalized voxel model, indicating that the RVM (after kernel regression) built from the QC’d experiments in these divisions is more capable of predicting regionalized connectivity calculated from held-out experiments within these regions.

However, we also note that, for the remaining 5/12 major divisions with excluded experiments, the retrained RVM and HM show higher model errors. This increase in error is particularly strong in the HM; for example, the HM error within the pallidum increases by 27.2%, and increases by 27% in the hypothalamus. Moreover, only 4 major divisions show improvements in the RVM and HM’s “power to predict” (PTP), defined as the regional error for held-out experiments with at least one other injection in that region which was not used for fitting. Still, we show that our QC process was particularly effective for hippocampal connectivity, showing improvements in voxel model error (-12.0%), RVM and HM error, and PTP.

### Graph Theoretical Analysis Reveals Subtle Organizational Changes Following QC

Following QC, we observed that the rich club coefficients and the topological rich club nodes remain mostly stable (Figure 5A and 5B). We observed the biggest loss in the rich club coefficient for the induseum griseum, a region for which our QC removed spurious connections to the medulla and other hindbrain regions (Figure 5A). We also observed that all ten nodes with the largest changes in rich club coefficient show a loss in the coefficient, including the induseum griseum of the hippocampus, as well as other hypothalamic, cortical, and striatal regions, including the well-connected caudoputamen. Additionally, we observed that all but three topological rich club nodes were preserved after QC (Figure 5B). The preserved regions showed similar distributions within the hippocampus, thalamus, and midbrain as a previous voxel-level analysis of the Knox et al. (2018) connectome (Coletta et al., 2020).

**Figure 5:**
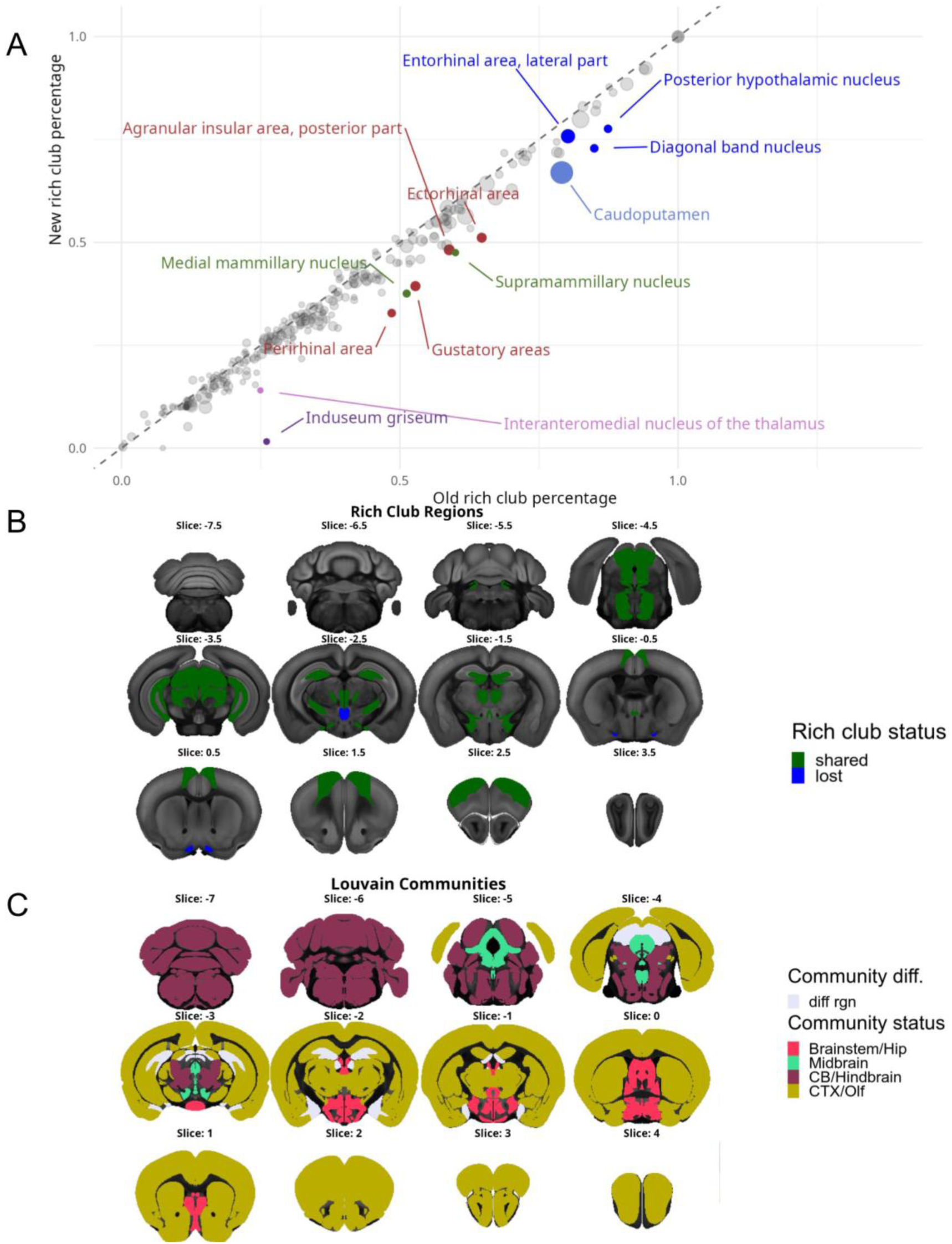
Graph theoretical analysis revealed organizational changes within the top 20% of connections in RVM, assuming symmetric connections across hemispheres. We observed changes in (A) the rich club coefficient, defined as the proportion of “rich” degrees that a given region’s degree is greater than or equal to (B) whether a region is “rich” or not, using a threshold of the mean plus one standard deviation of the “rich” degrees, and (C) the Louvain community assignments for each region.

After running the Louvain algorithm, we identified four communities in the old and new connectomes, which roughly correspond to brainstem/hippocampus, midbrain, cerebellum/hindbrain, and cortex/olfactory networks (Figure 5C). We further observed changes in the community assignments for five regions. We see one-off changes in community assignments for the lateral habenula of the thalamus, which changed from the cortical to the brainstem community, and the superior colliculus (motor-related), which changed from the midbrain to the CB/Hindbrain community. However, we also note that the dentate gyrus, posterior amygdalar nucleus, and medial amygdalar nucleus changed from the brainstem/hippocampus to the cortex/olfactory community.

To confirm that our graph theoretic results were not significantly biased by the size of the target region, we repeated our rich club and community detection analyses on the normalized connection density, which divides the normalized connection strength between two regions by the size of the target region. We observed similar rich club regions to before, as well as similarities in the rich club coefficient before and after QC, but with a few rich club regions gained after QC (Supplementary Figure 9A/B).

Additionally, we identified the same four communities as before, with four regions that changed community assignments. Of these four regions, three are thalamic nuclei (nucleus of reuniens, lateral habenula, paraventricular nucleus of the thalamus) that changed assignments from the cortical to the brainstem communities, and the remaining region (preparasubthalamic nucleus) changed assignments from the midbrain to the cortical community. Taken together, these results indicate subtle, metric-dependent organizational changes following QC.

## Discussion

Our quality control procedure identified a set of n=381 experiments that satisfy the credit assignment problem for building whole-brain connectomes; namely, projections were properly aligned and neither too small nor too diffuse to meaningfully represent axonal tracts, and injections were confined to a single brain region. After rebuilding the connectomes on this cleaner subset of the data, we observed large decreases in connection strength percentiles across major divisions. These changes arise from changes in regional-level connectivity, and suggest even stronger changes in voxel-level connectivity, which are particularly relevant to researchers examining specific connections to a single voxel or region of interest.

We excluded n=56 (∼13%) of the original 437 experiments, representing a variety of experimental and/or image processing failures. Additionally, of the n=17 experiments removed through our automated QC, n=11 of them were also identified during manual QC, confirming our initial automated filtering step (Supplementary Figure 1). Our automated QC using the brain mask identified three experiments that were grossly misaligned to the CCFv3 template, which could be due to a failure in the registration of the 2D microscopy images to the 3D template. The procedure used for this alignment involved stacking the 2D slices, assuming no missing slices during data collection (Kuan et al., 2015); however, inspecting the data mask for experiment 113036264 reveals missing coronal slices, and these missing slices appeared to not have been padded into the resulting 3D volume. This error resulted in a biologically invalid mask, driving the downstream registration misalignment. We note that, unlike the injection and projection failure modes, re-processing of the misaligned experiments could recover them for future use. We note that since the two-dimensional microscopy images were unavailable to download, we were unable to directly verify missing slices or alignment failures. However, we argue that our choice to manually QC all the data allowed us to detect the most egregious cases of misalignment, and our automated QC of out-of-brain voxels caught all but one of the misaligned experiments.

We also observed experiments that are upper and lower extrema for injection and projection voxel counts, or injections that span multiple major brain divisions. When algorithmically assigned to a single voxel or brain region, these experiments yield biologically unrealistic connectivity patterns, so we excluded these extreme cases to avoid over or under-representing connectivity to injected brain regions or centroid voxels. The large injections could have arisen from the electrical parameters of the iontophoresis technique used to control the virus’ entry into the injection site. The current strength and duration were chosen to produce infection areas of 400 to 700 µm, but it was reported that these parameters would result in larger infections (>1000 µm) in subcortical areas with lower neuron densities (Harris et al., 2012). These large injections could have also been due to a large or broken pipet tip (Harris et al., 2012).

We show that these large injections are highly correlated with larger projection volumes (Figure 2B). On the other hand, our manual and automated QC identify experiments with small injection sites and barely any projections beyond these injection sites, which could have been due to low currents or low titrated virus concentrations. For more information on experimental troubleshooting, see Table 2 of Harris et al. (2012).

Finally, our manual QC identified n=11 injections with cortical leakage, a failure mode likely caused by the passage of a leaking needle through the cortex when injecting deep, subcortical regions. Previous versions of the homogeneous and RVM assigned the projections from these experiments to both the primary and the secondary injection sites, or assigned the projection to the centroid of the overall injection volume (Oh et al., 2014; Knox et al., 2018). However, due to the difficulty in disentangling which projections were connected to the primary versus secondary injection site, and since the injection centroid’s location was biased by the off-target cortical injection volume, we chose to remove these experiments.

After our QC, we rebuilt the homogeneous and RVMs, keeping all algorithmic parameters the same but removing the n=56 failed experiments. Notably, in both models, we observed strong continuous losses and gains in connectivity aggregated across major brain divisions, with stronger losses and gains in contralateral than ipsilateral connectivity (Figure 4). The losses in contralateral connectivity are likely due to the n=8 experiments that were removed due to nonspecific tracer spreading, whereas the gains in connectivity are due to removal of n=15 experiments with small amounts of tracer spreading from the injection site, which could have originally diluted contralateral connectivity to a given brain region. Since the RVM averages distance-weighted projections across injected major divisions, it is likely to produce gains in contralateral connectivity in divisions with small, removed injections or projections (ex: olfactory bulb) and losses in connectivity to regions with large, removed injections or projections (ex: hippocampus) (Supplementary Figure 5). Similarly, since the HM fits a linear model to predict whole-brain projection data from injection data in a brain region, it is also likely to gain contralateral connectivity in regions where small projections are removed, and lose contralateral connectivity in regions where large projections are removed, as seen in the olfactory bulb and hippocampus (Supplementary Figure 6).

We interpreted our post-QC percentile changes (Supplementary Figures 5A and 6A) as losses in false-positive connections and gains in true-positive connections. Follow-up analyses, solely examining the top 20% of connection strengths in the old versus new connectomes, revealed losses in cerebellum-isocortex and hippocampus-medulla connections in both models. To our knowledge, previous tractography studies in mice do not show monosynaptic connections between the cerebellum and isocortex (Schröder et al., 2020; Kang et al., 2021) or between the hippocampus and medulla (Schröder et al., 2020; Cembrowski et al., 2018; Qiu et al., 2024); therefore, we view these connections as implausible. As a result of downweighting these implausible connections, in both models, we also observed gains in previously-characterized hypothalamus-isocortex (Jiao et al., 2025; Risold et al., 1997) connectivity specifically in the lateral zone of the hypothalamus, hypothalamus-cerebellum connectivity (Onat and Çavdar 2003; Zhu et al., 2006; Dietrichs 1984), and midbrain-cerebellum connectivity, including connections between two raphe nuclei and the paraflocculus of the cerebellum in the RVM (Oostland and Hooft 2016; Waterhouse et al., 1986), further confirming our approach. When examining the top 5% of connections, the 100% loss in hippocampus-medulla connections in the RVM persisted; however, the other changes in connectivity only existed at the 20% threshold, underscoring the need to explore which connectomic changes persist across a variety of connectome density thresholds.

Despite these consistencies, we also observed differences in the changes in both connectomes after QC due to the inherent algorithmic differences between the RVM and HM. These differences include the assumption of “smooth” connectivity patterns within a single major brain division in the RVM due to the Gaussian kernel, meaning that even if an injected region loses connectivity after QC, its connectivity can still be approximated by the spatial average of the nearby injection sites. In the HM, there is no averaging of experiments across regions; hence, if a region loses a large proportion of injections after QC, there is less projection data used to predict its whole-brain connectivity through the nonnegative linear regression, resulting in the potential assignment of spurious connectivity patterns. For example, we see large losses in connectivity to an olfactory region (bottom right slice of Figure 3B), and the RVM and HM respond differently to this loss. Strikingly, we see a 100% loss in the top 20% of connections between the olfactory bulb and ipsilateral medulla in the RVM (Supplementary Figure 5B) but a 100% gain in the HM (Supplementary Figure 6B), as well as a 100% loss in contralateral striatum-medulla connections in the RVM, but a 500% gain in the HM. Therefore, to ensure robustness to missing data, we suggest using the RVM over the HM, since the distance-weighted average of connectivity within broad major brain divisions is inherently more robust to missing data, rather than the HM’s assignment of connectivity to only the brain regions immediately injected. Indeed, due to the lack of existing evidence for monosynaptic olfactory bulb-medulla or striatum-medulla connections (Imamura et al., 2020; Schröder et al., 2020; Märtin et al., 2019; Nakano et al., 2000), we posit that the RVM is a more biologically accurate representation of the mouse structural connectome.

Finally, as a vignette to demonstrate architectural changes following QC, we repeated a modified version of the rich club and community detection algorithm with a binarized matrix of the top 20% of connections in the RVM (Figure 5) (Herrera-Portillo et al., 2025; Coletta et al., 2020). Notably, our QC’d, regionalized connectome showed consistencies with previous graph theoretical analyses at the voxel level (Coletta et al., 2020), and our QC process minimally changed the overall rich club regions. Consistent with Coletta et al. (2020), we implicate the nucleus reuniens of the thalamus as a rich club region, which has been shown to affect cognition in perturbational studies in mice (Vetere et al. 2017). We also showed the emergence of four communities in our structural connectome, with subtle changes in community assignments for n=6 regions after QC. Certain thalamic regions, including the lateral habenula, switched assignments from cortical to brainstem/hippocampus communities, consistent with previous tracer experiments showing direct habenula-cerebellum and anteromedial nucleus-hippocampal connections (Huang et al., 2024; de Lima et al., 2017). Additionally, prior work in mice has implicated a circuit connecting the olfactory bulb, amygdala, and hippocampus, which is recapitulated in our changed community assignment for the dentate gyrus, posterior and medial amygdalar nuclei to the cortex/olfactory community after QC (Cadiz-Moretti et al., 2016; Schröder et al., 2020). Nonetheless, this community reorganization differs after normalizing connection strengths by region size (Supplementary Figure 9B) (i.e. normalized connection density), necessitating further analysis using isotropic voxels, or equally-sized parcellations. We also note that the rich club and community metrics are biased by the most highly connected brain regions, which did not change after QC. We encourage future work to explore the effects of our QC’d connectomes on metrics that are more sensitive to connectomic changes; for example, the “average shortest path” metric is biased by adding a single false positive or false negative connection, resulting in many nodes with significantly shortened paths (Kitsak et al., 2023)

Although we interpret our rebuilt connectomes as more biologically plausible than the existing connectomes, our work has several limitations. Firstly, our QC depends on binary thresholding. We justified this choice based on the exclusion of noisy background fluorescence, but our thresholding choice could also artificially exclude meaningful connections. Nonetheless, one of our raters (V.N.) repeated the manual QC for all injection and projection data at a lower (0.01) threshold and saw similar results (Supplementary Data 1), suggesting that our failures were not threshold-dependent.

Additionally, we did not change the algorithms underlying the RVM or HM to accommodate for the lower number of included experiments, which could result in the overfitting of connectivity to certain brain regions with large losses in injection experiments, such as the cerebellar or hypothalamic regions with a complete loss in injection experiments (Figure 3B). Thirdly, we cannot exclude the possibility that experiments with the smallest projection volumes, especially in the cerebellum, meaningfully represent short-range connectivity to smaller axons, such as cerebellar Purkinje cell axons (Perge et al., 2012); however, due to their small volumes, these experiments have limited utility in whole-brain contexts like MRI-based analyses (Yee et al., 2018), and are similar to previously-excluded experiments in Knox et al. (2018), confirming our decision to exclude them. Finally, for experiments with off-target injections, we discarded the entire experiment, resulting in a loss in connectivity.

However, a more sophisticated approach could disentangle the connectivity to voxels in the intended target injection region, resulting in the inclusion of more experimental data.

Despite the improvements uncovered by our QC, an inherent limitation to all AMBCA-derived murine connectomes is incomplete sampling. Firstly, these connectomes were only built from data from male C57BL/6J mice, reducing generalizability across sexes and strains based on known sex and strain effects on structural connectivity patterns (Elkind et al., 2023; Karatas et al., 2021). Additionally, we observe major divisions with very few experiments, such as the pallidum with 10 experiments and the pons with 16 experiments. Such limited data may not be sufficient to properly characterize connectivity within these regions, and may also be insufficient to account for the inter-individual variability that is known to exist in the murine structural connectome (Lopes et al., 2023). Nonetheless, the AMBCA also contains many cell-type-specific connectivity experiments (n=2,995 total experiments) many of which use Cre driver lines on a C57BL/6J genetic background. Since these connections are, by definition, a subset of wild-type connectivity patterns, they can be integrated into the RVM and HM algorithms to improve the spatial coverage and reliability of the estimated connectomes.

Additionally, since the inception of the AMBCA, there have been numerous additional efforts to characterize the murine whole-brain connectome using computational single-cell reconstructions (Xiong et al., 2025), additional two-photon microscopy experiments (Winnubst et al., 2019), fluorescent light-sheet microscopy (Son et al., 2022), and ex-vivo magnetic resonance imaging (Arefin et al., 2021; Crater et al., 2022; Allan Johnson et al., 2023). These modalities could serve as independent validations for the connections in our QC’d RVM and HM connectomes, or could be integrated into future, multimodal models of connectivity.

The AMBCA is the product of an immense scientific effort by the Allen Institute, and has been incredibly valuable to the neuroscientific community. To further improve its use in light of our findings, we encourage the community to use our rebuilt connectomes, leveraging the data within a subset of the AMBCA to produce more reliable estimates of structural connectivity in the mouse brain. We also have provided evidence for the fact that our rebuilt connectomes subtly alter downstream analyses applying whole-brain structural connectivity, including the characterization of rich clubs and communities at the regional level. We anticipate that our QC’d connectomes will have downstream impacts on a wide variety of mechanistic and pathological models using the original connectomes (Tullo et al., 2025; Rahayel et al., 2022; Henderson et al., 2019; Lubben et al., 2024). Moreover, since the AMBCA experiments were used to build the CCFv3 template, future work needs to assess the effects of excluding misaligned experiments on the anatomy of this commonly-used template. Finally, we hope that future work will extend the application of our QC criteria to the entire AMBCA, consisting of n=2,995 experiments in C57BL/6 mice and other transgenic mouse lines that visualize specific cell types of interest.

## Supplementary Data

Our rebuilt connectomes and QC images are publicly available at the following Zenodo repository: https://zenodo.org/records/20276701.

## Data and Code Availability

All code is stored on https://github.com/vik16nathan/allen_connectome_qc. All input data is publicly available and downloaded from the Allen API.

## Ethics

All data in this work is downloaded from the publicy-available Allen Mouse Brain Connectivity Atlas (AMBCA) (Oh et al., 2014). As mentioned in the reference publication, “All experimental procedures related to the use of mice were approved by the Institutional Animal Care and Use Committee of the Allen Institute for Brain Science, in accordance with NIH guidelines.”

## Author Contributions

V.N.: Conceptualization, Software, Formal Analysis, Investigation, Writing—Original Draft, Writing—Review & Editing. S.T.: Formal Analysis, Writing—Review & Editing. L.H.: Formal Analysis, Writing—Review & Editing. G.D.: Conceptualization, Methodology, Writing—Review & Editing. Y.Y.: Conceptualization, Methodology, Writing—Review & Editing. M.M.C.: Conceptualization, Supervision, Writing—Review & Editing, Funding Acquisition.

## Acknowledgements

We would like to thank Kameron Decker Harris, Jennifer Whitesell, and Stefan Mihalas at the Allen Institute for thoughtful discussions about the tracer data processing involved in building the connectome from Knox et al. (2018). We would also like to thank the authors of Knox et al. (2018), Joseph E. Knox, Kameron Decker Harris, Nile Graddis, Jennifer Whitesell, Hongkui Zeng, Julie A. Harris, Eric Shea-Brown and Stefan Mihalas, for providing the code to recreate the RVM and HM connectomes.

## Supplementary Tables

**Supplementary Table 1:** List of abbreviations for major division names.

| Full Major Division Name | Major Division Abbreviation |
| --- | --- |
| Isocortex | Isocortex |
| Olfactory areas | OLF |
| Hippocampal formation | HPF |
| Cortical subplate | CTXsp |
| Striatum | STR |
| Pallidum | PAL |
| Thalamus | Thal |
| Hypothalamus | Hypothal |
| Midbrain | Midbrain |
| Pons | Pons |
| Medulla | Medulla |
| Cerebellum | CB |

**Supplementary Table 2:** The number of experiments with injections in each major brain division before versus after QC.

| Division | Before QC | After QC | Lost |
| --- | --- | --- | --- |
| Isocortex | 128 | 121 | 7 |
| OLF | 21 | 19 | 2 |
| HPF | 49 | 39 | 10 |
| CTXsp | 8 | 7 | 1 |
| STR | 27 | 23 | 4 |
| PAL | 10 | 9 | 1 |
| Thal | 44 | 38 | 6 |
| Hypothal | 34 | 25 | 9 |
| Midbrain | 42 | 33 | 9 |
| Pons | 16 | 16 | 0 |
| Medulla | 39 | 35 | 4 |
| CB | 19 | 16 | 3 |

**Supplementary Table 3:**
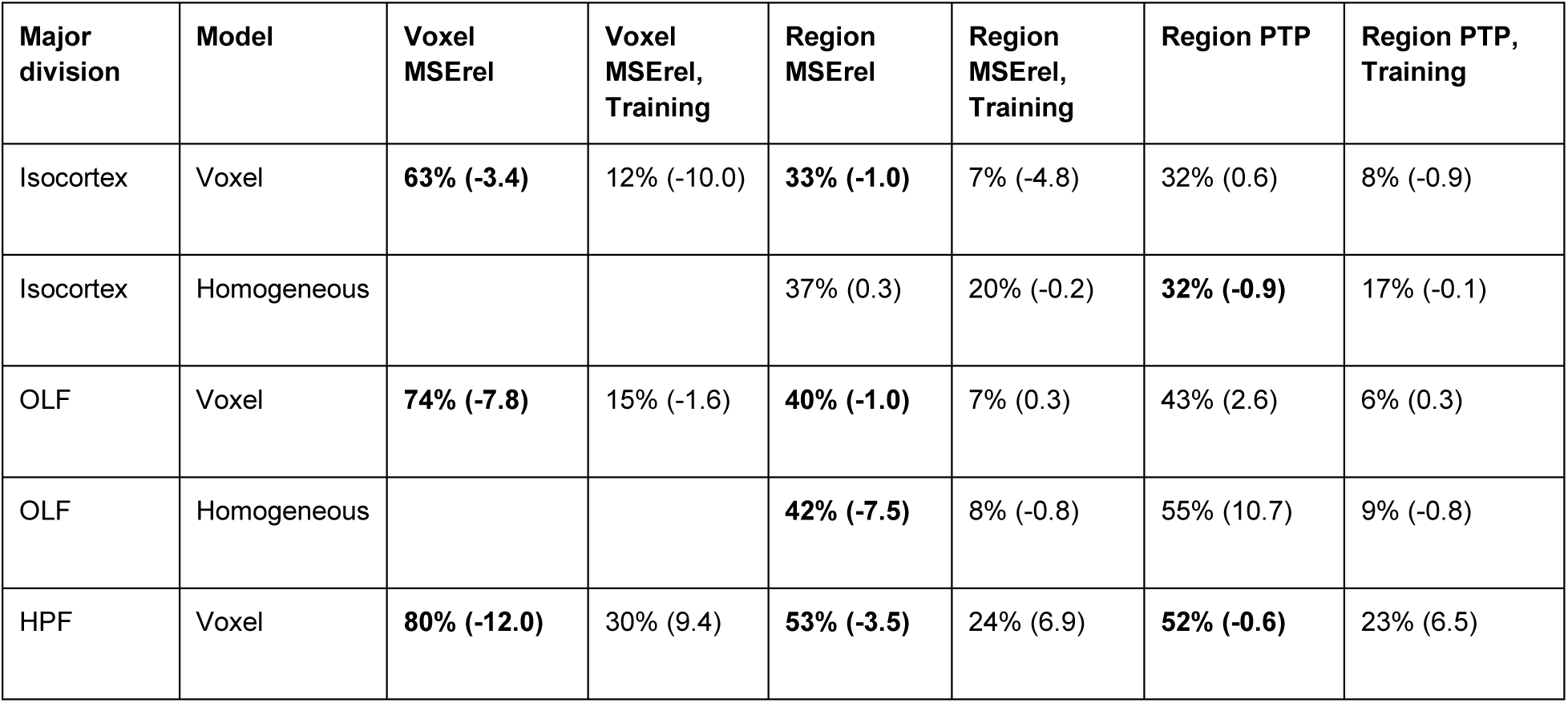
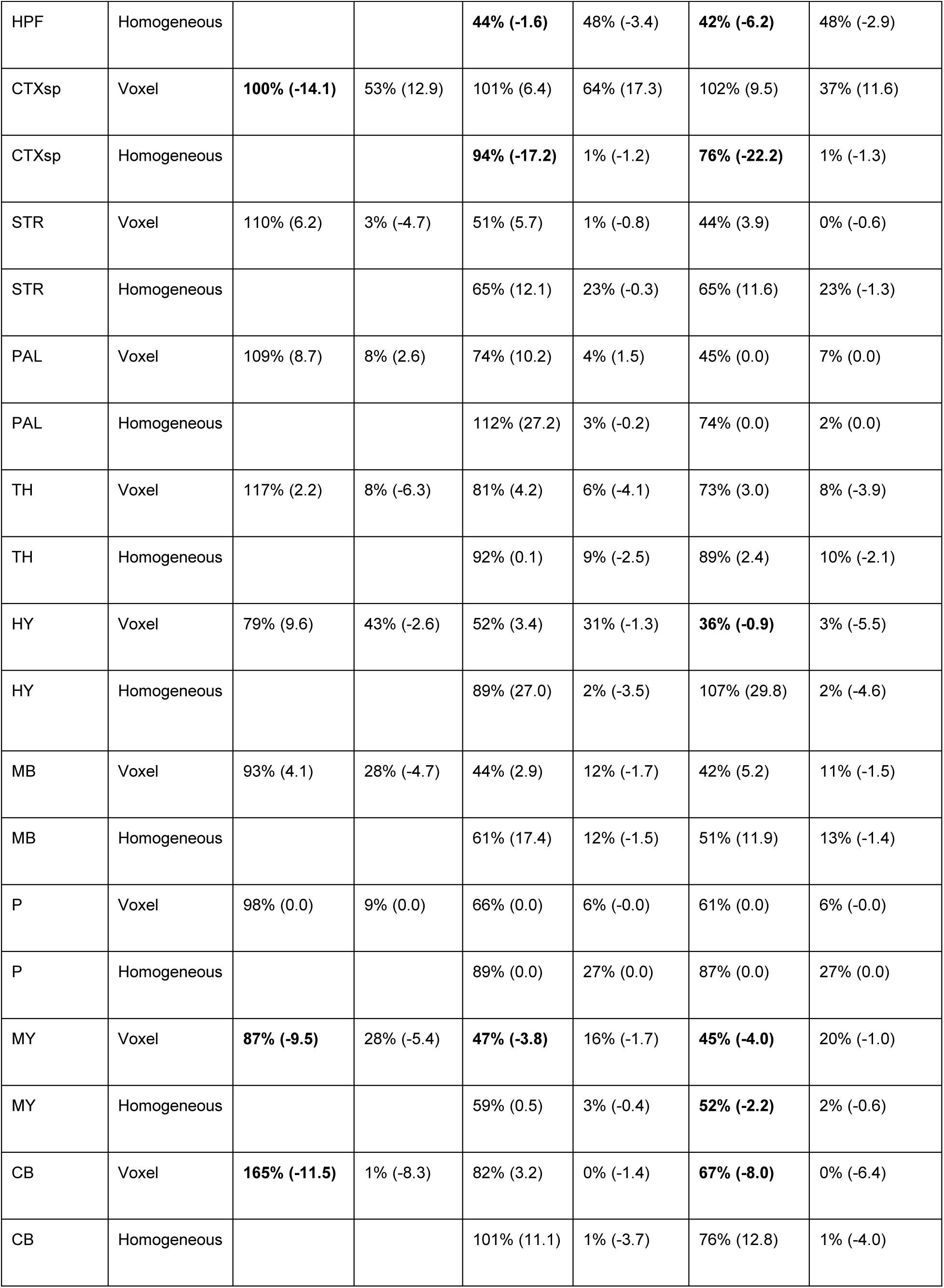
We replicate Table 1 from Knox et al., which calculates MSE_rel_ values, defined as “the mean square error relative to the average squared norm of the prediction and left-out data.” These values can range from 0-200% (Knox et al., 2018). In parentheses, we also show the change in error after QC. Bolded values indicate reductions in error after QC versus before QC.

| Major division | Model | Voxel $MSE_{rel}$ | Voxel $MSE_{rel}$, Training | Region $MSE_{rel}$ | Region $MSE_{rel}$, Training | Region PTP | Region PTP, Training |
| --- | --- | --- | --- | --- | --- | --- | --- |
| Isocortex | Voxel | **63% (-3.4)** | 12% (-10.0) | **33% (-1.0)** | 7% (-4.8) | 32% (0.6) | 8% (-0.9) |
| Isocortex | Homogeneous |  |  | 37% (0.3) | 20% (-0.2) | **32% (-0.9)** | 17% (-0.1) |
| OLF | Voxel | **74% (-7.8)** | 15% (-1.6) | **40% (-1.0)** | 7% (0.3) | 43% (2.6) | 6% (0.3) |
| OLF | Homogeneous |  |  | **42% (-7.5)** | 8% (-0.8) | 55% (10.7) | 9% (-0.8) |
| HPF | Voxel | **80% (-12.0)** | 30% (9.4) | **53% (-3.5)** | 24% (6.9) | **52% (-0.6)** | 23% (6.5) |
| HPF | Homogeneous |  |  | **44% (-1.6)** | 48% (-3.4) | **42% (-6.2)** | 48% (-2.9) |
| CTXsp | Voxel | **100% (-14.1)** | 53% (12.9) | 101% (6.4) | 64% (17.3) | 102% (9.5) | 37% (11.6) |
| CTXsp | Homogeneous |  |  | **94% (-17.2)** | 1% (-1.2) | **76% (-22.2)** | 1% (-1.3) |
| STR | Voxel | 110% (6.2) | 3% (-4.7) | 51% (5.7) | 1% (-0.8) | 44% (3.9) | 0% (-0.6) |
| STR | Homogeneous |  |  | 65% (12.1) | 23% (-0.3) | 65% (11.6) | 23% (-1.3) |
| PAL | Voxel | 109% (8.7) | 8% (2.6) | 74% (10.2) | 4% (1.5) | 45% (0.0) | 7% (0.0) |
| PAL | Homogeneous |  |  | 112% (27.2) | 3% (-0.2) | 74% (0.0) | 2% (0.0) |
| TH | Voxel | 117% (2.2) | 8% (-6.3) | 81% (4.2) | 6% (-4.1) | 73% (3.0) | 8% (-3.9) |
| TH | Homogeneous |  |  | 92% (0.1) | 9% (-2.5) | 89% (2.4) | 10% (-2.1) |
| HY | Voxel | 79% (9.6) | 43% (-2.6) | 52% (3.4) | 31% (-1.3) | **36% (-0.9)** | 3% (-5.5) |
| HY | Homogeneous |  |  | 89% (27.0) | 2% (-3.5) | 107% (29.8) | 2% (-4.6) |
| MB | Voxel | 93% (4.1) | 28% (-4.7) | 44% (2.9) | 12% (-1.7) | 42% (5.2) | 11% (-1.5) |
| MB | Homogeneous |  |  | 61% (17.4) | 12% (-1.5) | 51% (11.9) | 13% (-1.4) |
| P | Voxel | 98% (0.0) | 9% (0.0) | 66% (0.0) | 6% (-0.0) | 61% (0.0) | 6% (-0.0) |
| P | Homogeneous |  |  | 89% (0.0) | 27% (0.0) | 87% (0.0) | 27% (0.0) |
| MY | Voxel | **87% (-9.5)** | 28% (-5.4) | **47% (-3.8)** | 16% (-1.7) | **45% (-4.0)** | 20% (-1.0) |
| MY | Homogeneous |  |  | 59% (0.5) | 3% (-0.4) | **52% (-2.2)** | 2% (-0.6) |
| CB | Voxel | **165% (-11.5)** | 1% (-8.3) | 82% (3.2) | 0% (-1.4) | **67% (-8.0)** | 0% (-6.4) |
| CB | Homogeneous |  |  | 101% (11.1) | 1% (-3.7) | 76% (12.8) | 1% (-4.0) |

**Supplementary Table 4:** Agreement in ratings across QC criteria for injection and projection QC images amongst the two raters (VN and ST). Due to its rarity, we deliberately do not show “5” failures for projections, since there was only one experiment that VN rated with a 5.

|  | ST 0 | ST 1 | ST 2 | ST 4 |
| --- | --- | --- | --- | --- |
| VN 0 | 361 | 31 | 5 | 8 |
| VN 1 | 1 | 3 | 0 | 0 |
| VN 2 | 1 | 0 | 5 | 0 |
| VN 4 | 2 | 12 | 2 | 0 |

4B: Projection ratings
|  | ST 0 | ST 1 | ST 2 | ST 3 | ST 4 |
| --- | --- | --- | --- | --- | --- |
| VN 0 | 368 | 5 | 34 | 1 | 5 |
| VN 1 | 4 | 0 | 0 | 0 | 0 |
| VN 2 | 0 | 0 | 4 | 0 | 0 |
| VN 3 | 0 | 0 | 0 | 3 | 0 |
| VN 4 | 11 | 0 | 0 | 0 | 1 |

## Supplementary Figures

**Supplementary Figure 1:**
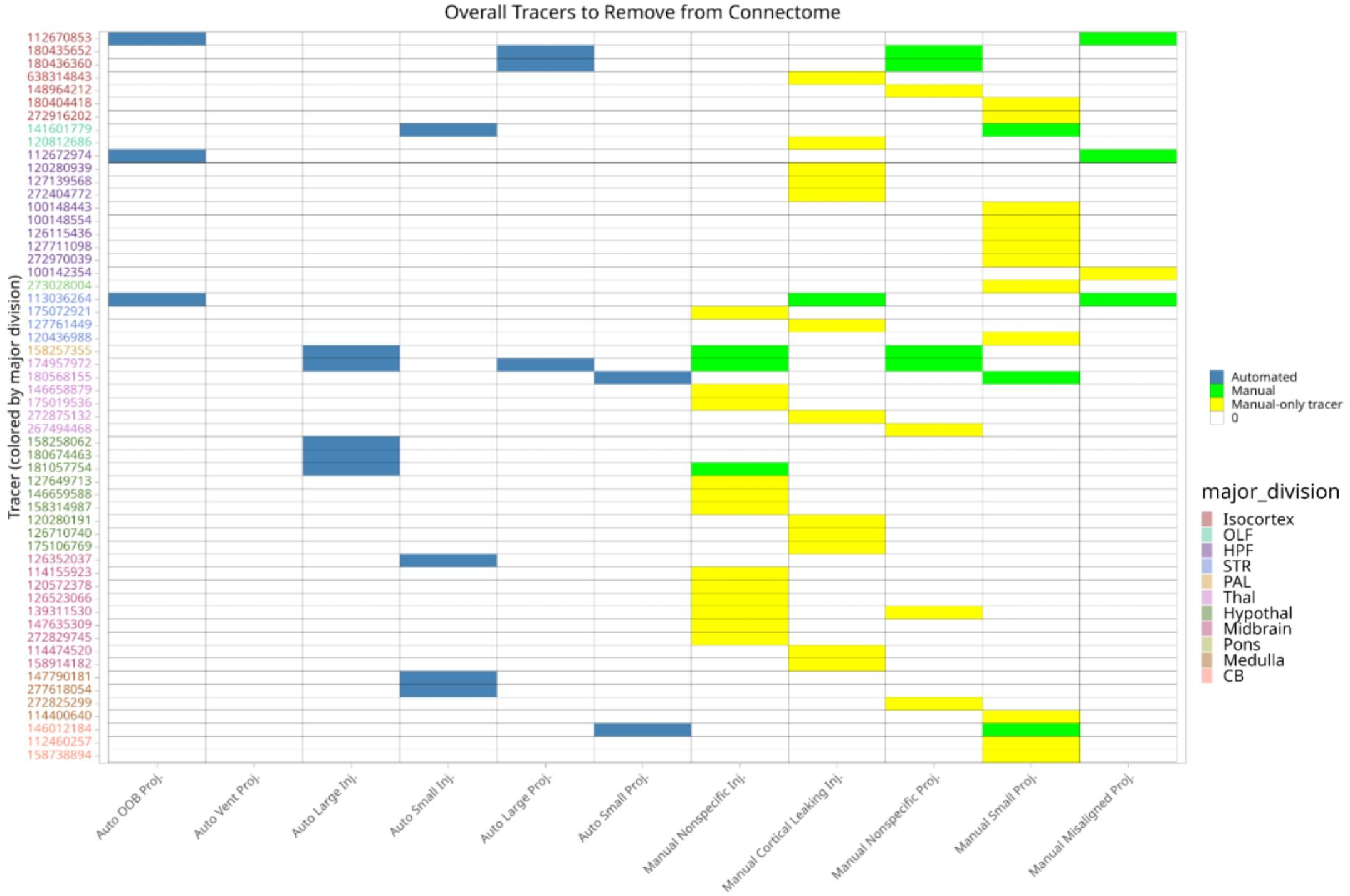
The complete set of n=56 experiments removed during our QC procedure, organized by the major division of the experimental injection. For each removed experiment, we show the failure mode(s) that result in the experiment’s exclusion, differentiating between automated removal criteria in blue and manual removal criteria in yellow/green. If an experiment has a green label for a manual removal criteria, then it was also flagged during automated QC.

**Supplementary Figure 2:**
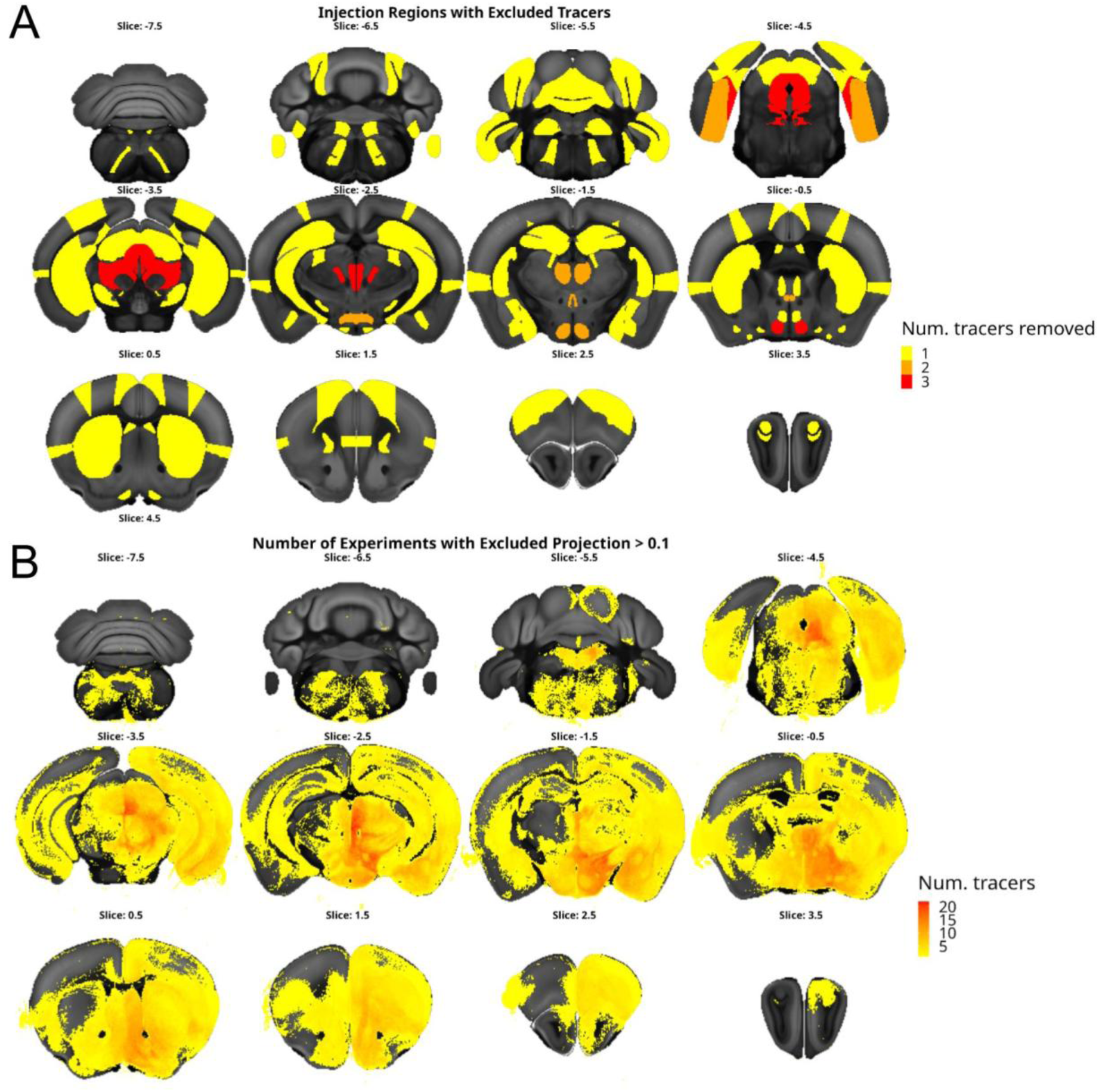
(A) The number of experimental injections removed from each brain region and (B) the number of experimental projections removed from each voxel, aggregated across all n=56 removed experiments. Note that we chose to show the injections removed per region rather than per voxel due to the relative sparsity of the injection sites compared to the projection data.

**Supplementary Figure 3:**
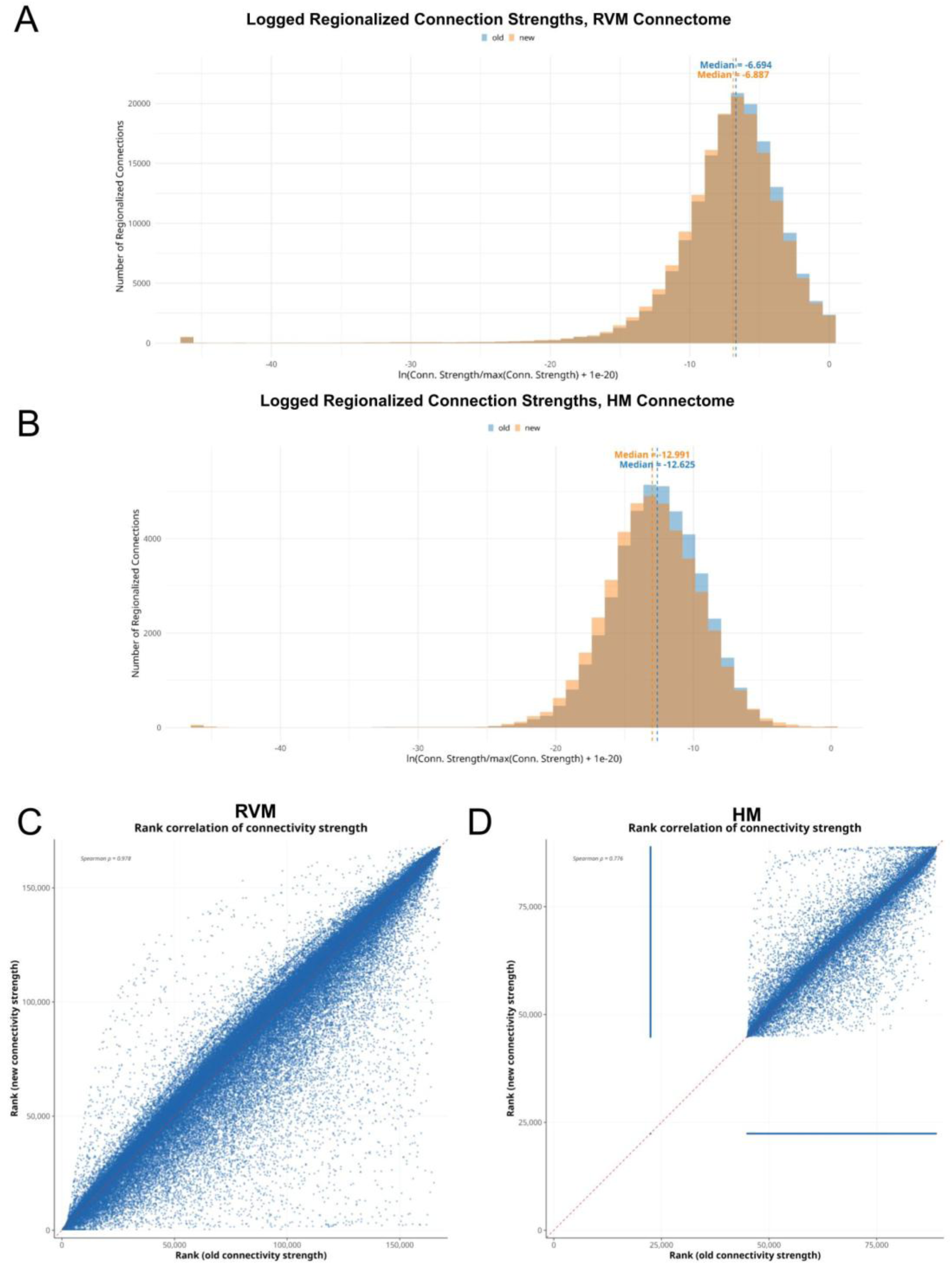
Histograms showing the distributions of the logged, normalized connection strengths in (A) the RVM and (B) the HM. In each panel, we compare the distribution of the connection strengths before QC (blue) versus after QC (orange).

**Supplementary Figure 4:**
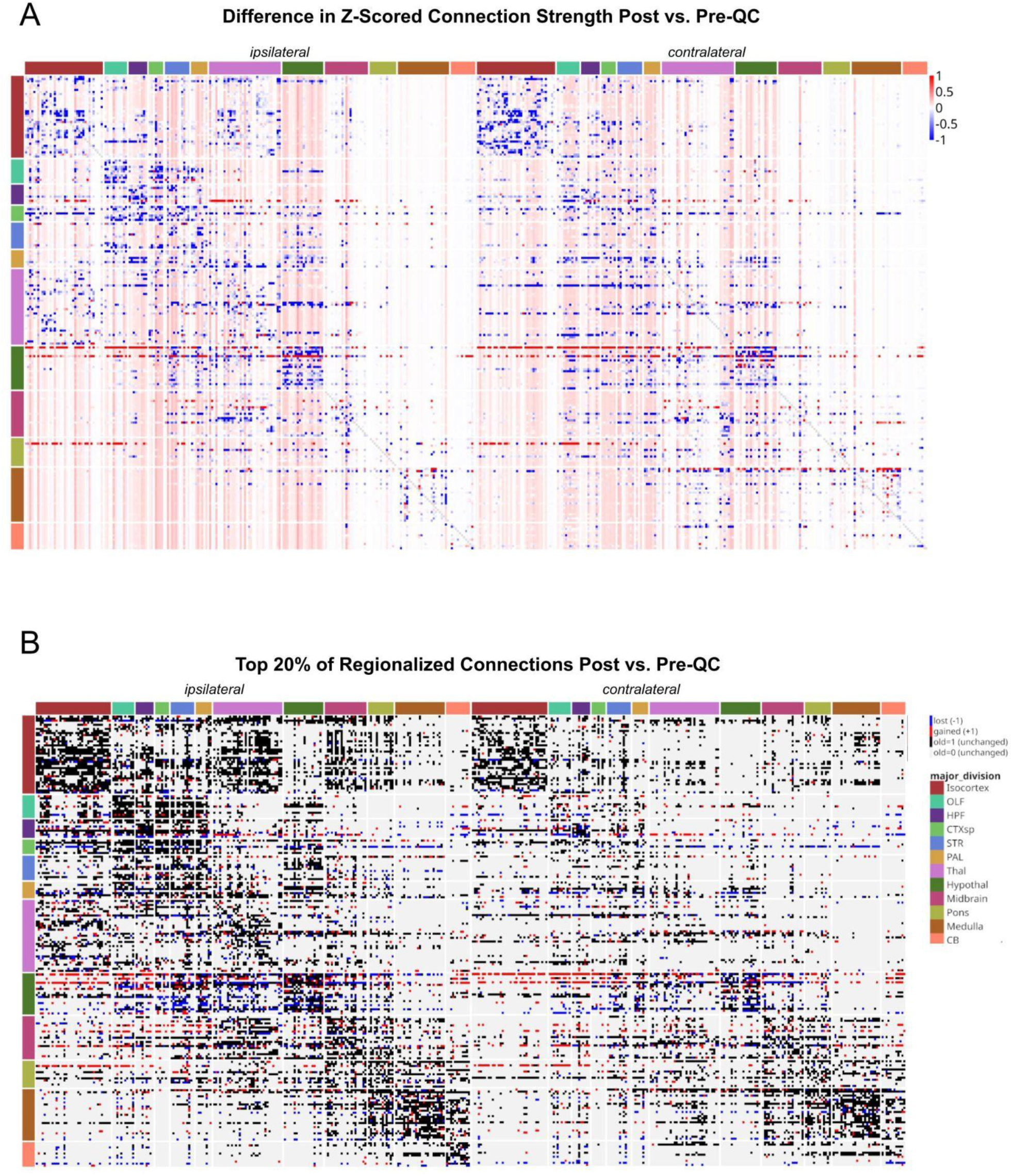
Similar to Figure 4, we observe whole-brain architectural changes in the HM (n=211 regions) after QC, when examined (A) continuously, when examining the difference in z-scored, normalized connection strengths, thresholded from [-1,1] due to the presence of outliers (B) discretely, when examining the lost (blue), gained (red) and unchanged (black/gray) connections within the subset of the top 20% of the old vs. rebuilt connectomes.

**Supplementary Figure 5:**
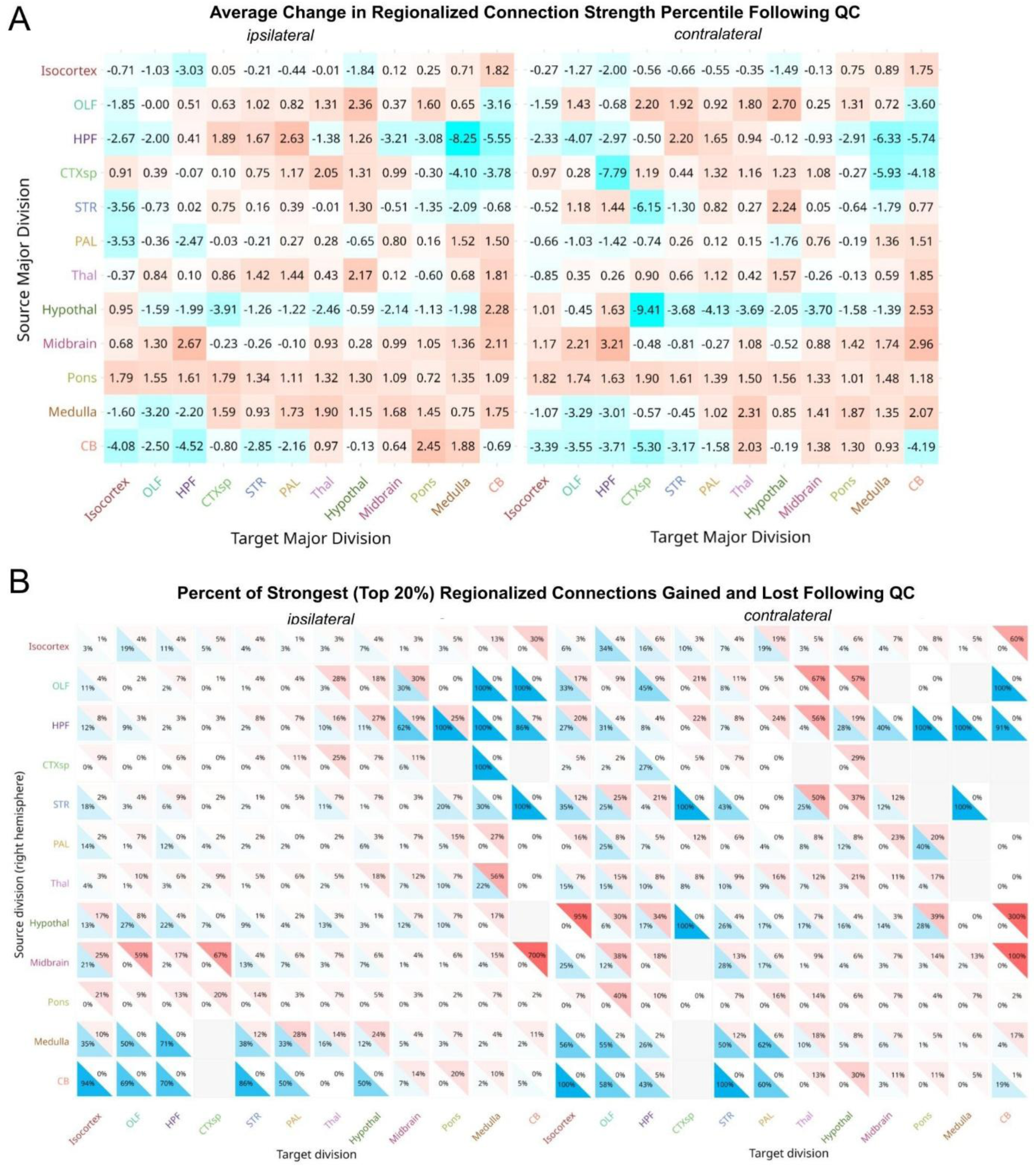
We aggregate the changes in connectivity from the RVM, showing (A) the average change in connection percentile across all connections and (B) the proportion of the top 20% of connections that were lost or gained, relative to the original number of connections in each major division in the RVM, summarized across each pair of major brain divisions from Knox et al.. For both plots, blue denotes a loss in connectivity and red denotes a gain in connectivity.

**Supplementary Figure 6:**
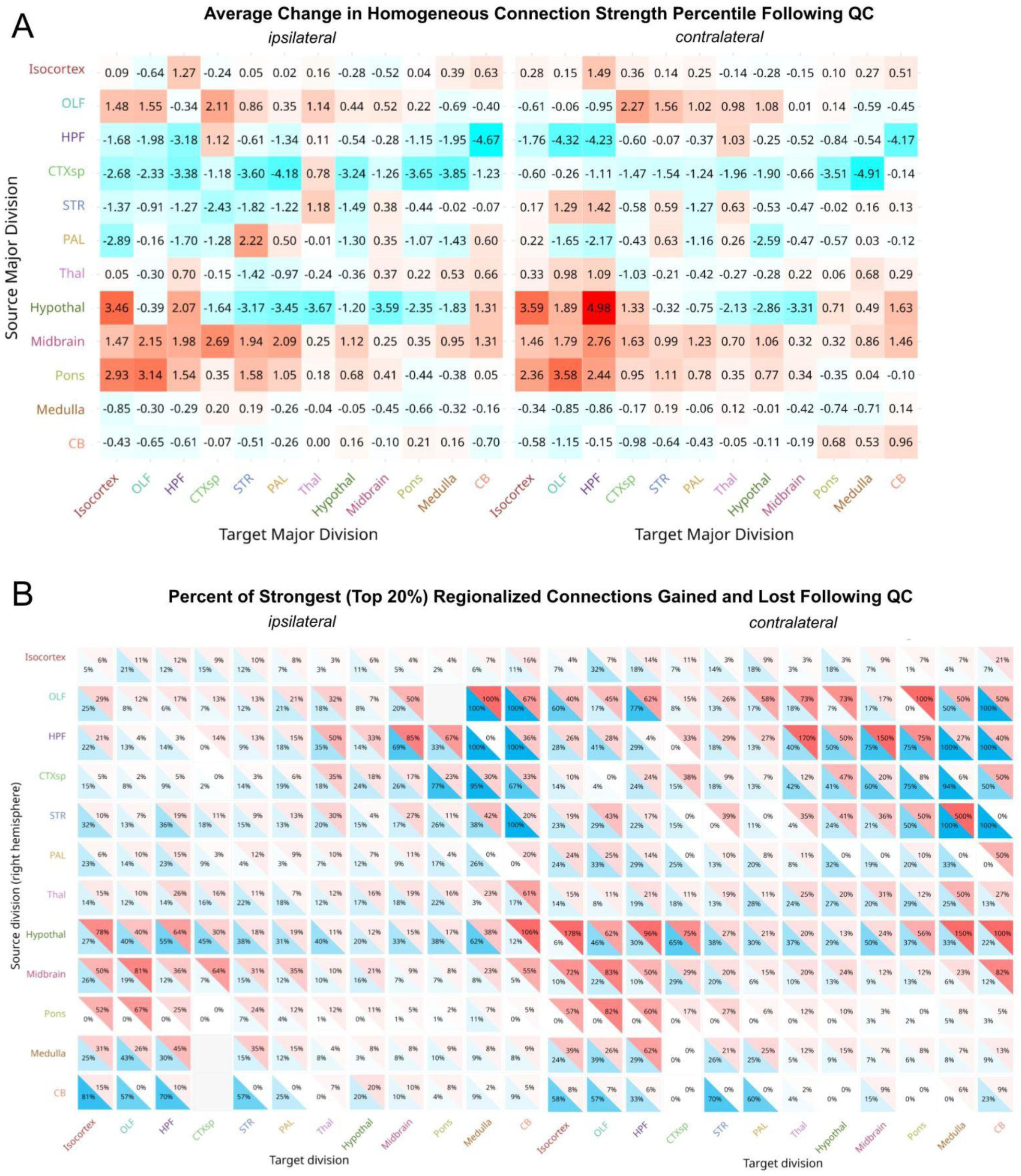
Similar to Supplementary Figure 5, we aggregate the changes in connectivity from the HM, showing (A) the average change in connection percentile across all connections and (B) the proportion of the top 20% of connections that were lost or gained, summarized across each pair of major brain divisions from Oh et al.. For both plots, blue denotes a loss in connectivity and red denotes a gain in connectivity.

**Supplementary Figure 7:**
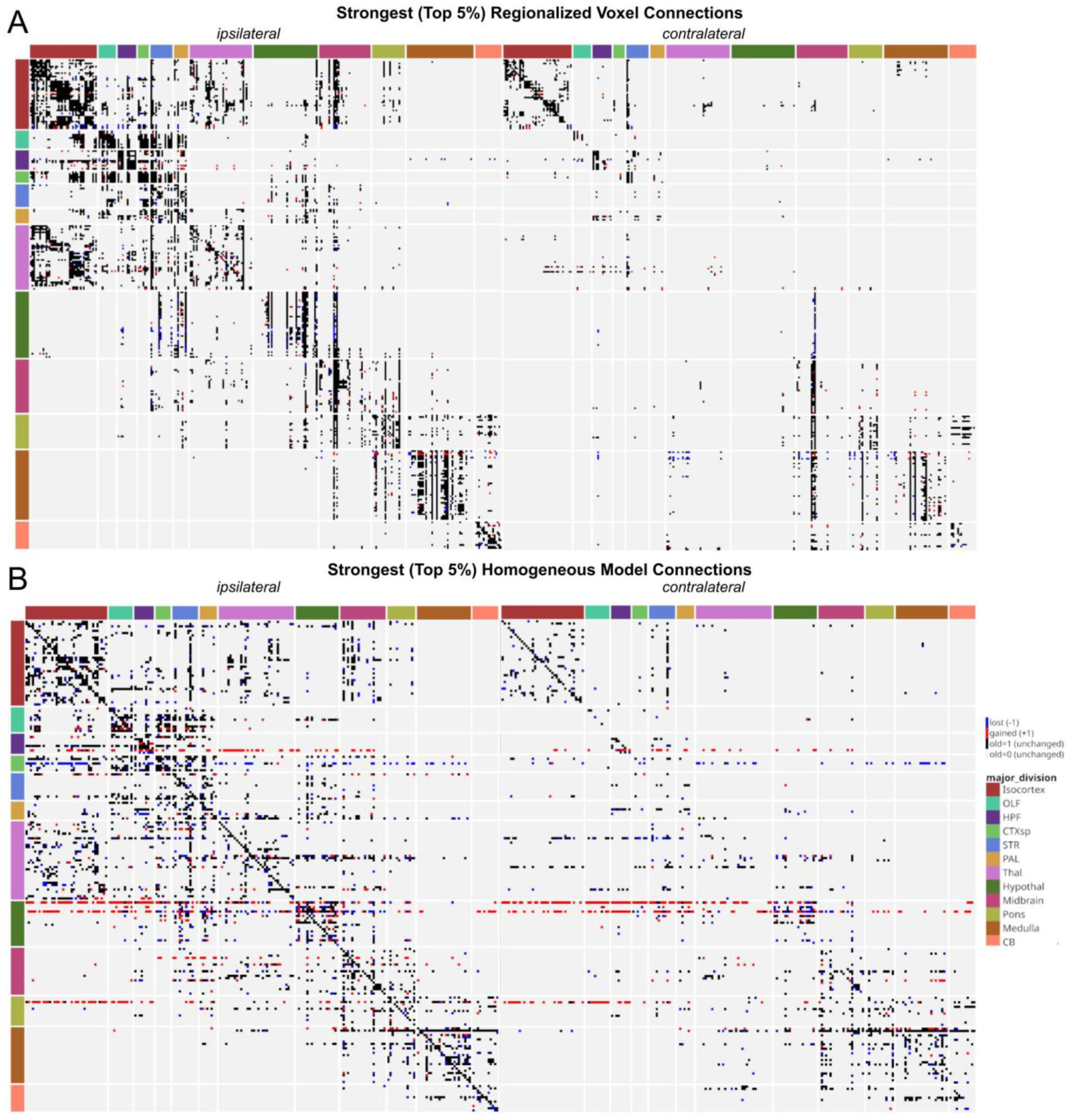
Similar to Figure 4B, we observe discrete connectivity changes when examining the top 5% of connection strengths in the old vs. rebuilt connectomes. We show lost (blue), gained (red) and unchanged (black/gray) connections within the (A) RVM (B) HM.

**Supplementary Figure 8:**
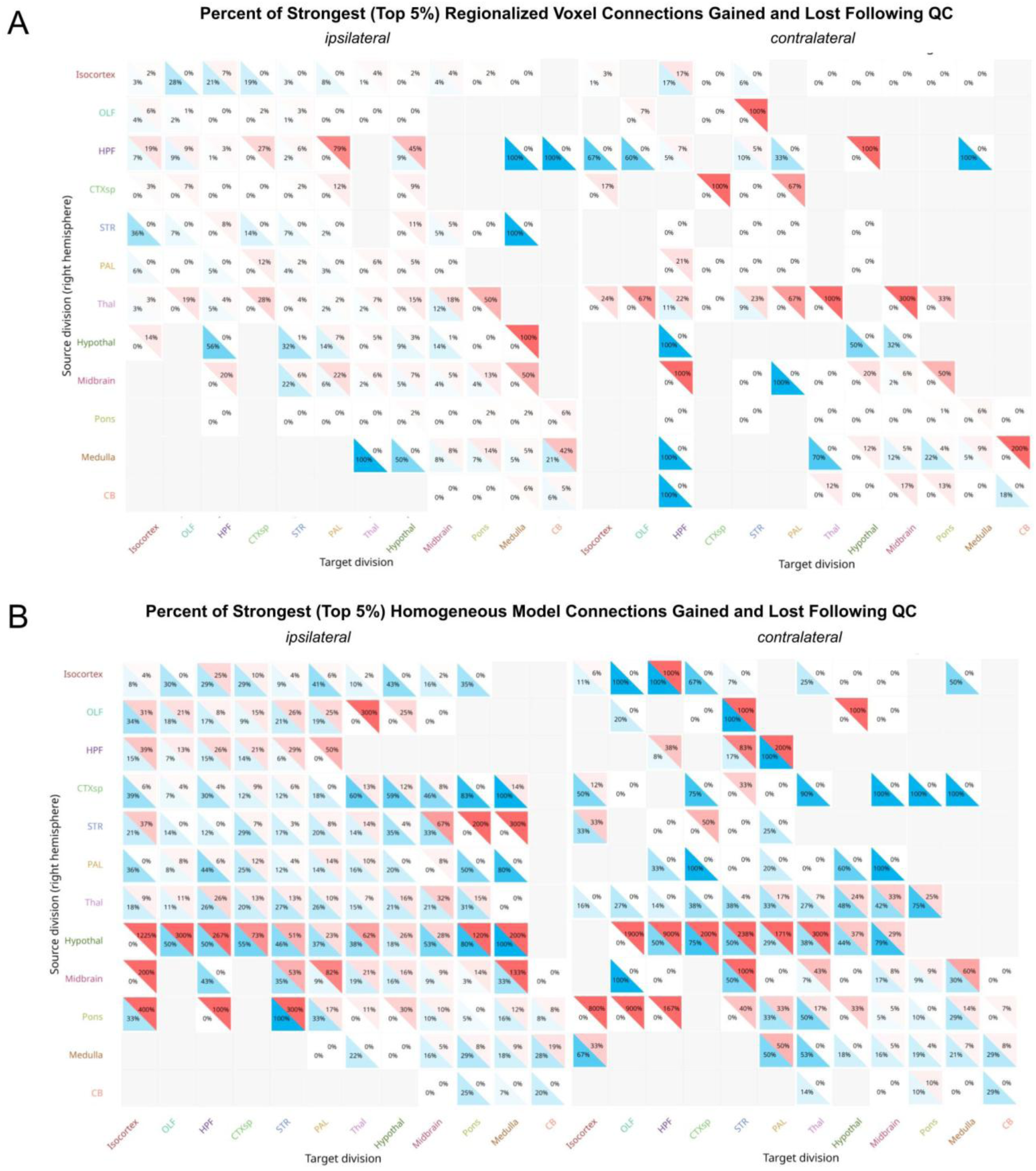
Similar to Supplementary Figure 5B, we show the proportion of the top 5% of connections that were lost or gained, summarized across each pair of major brain divisions in (A) the RVM and (B) the HM.

**Supplementary Figure 9:**
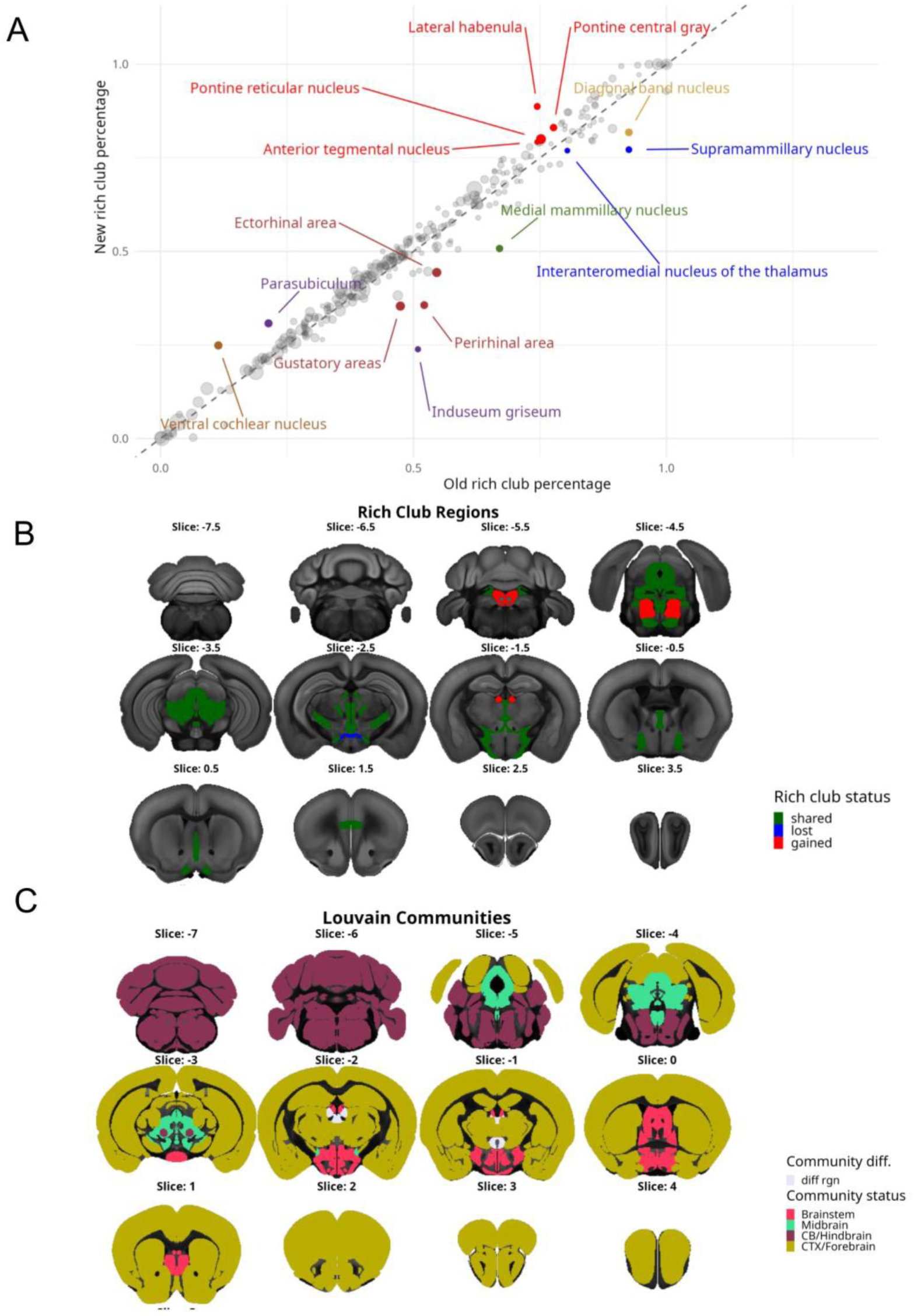
Similar to Figure 5, we show organizational changes within the top 20% of connections, using the “normalized connection density” instead of the normalized connection strength in the RVM. We assume symmetric connections across hemispheres and see changes in (A) the rich club coefficient, defined as the proportion of “rich” degrees that a given region’s degree is greater than or equal to (B) whether a region is “rich” or not, using a threshold of the mean plus one standard deviation of the “rich” degrees, and (C) the Louvain community assignments for each region.

**Supplementary Figure 10:**
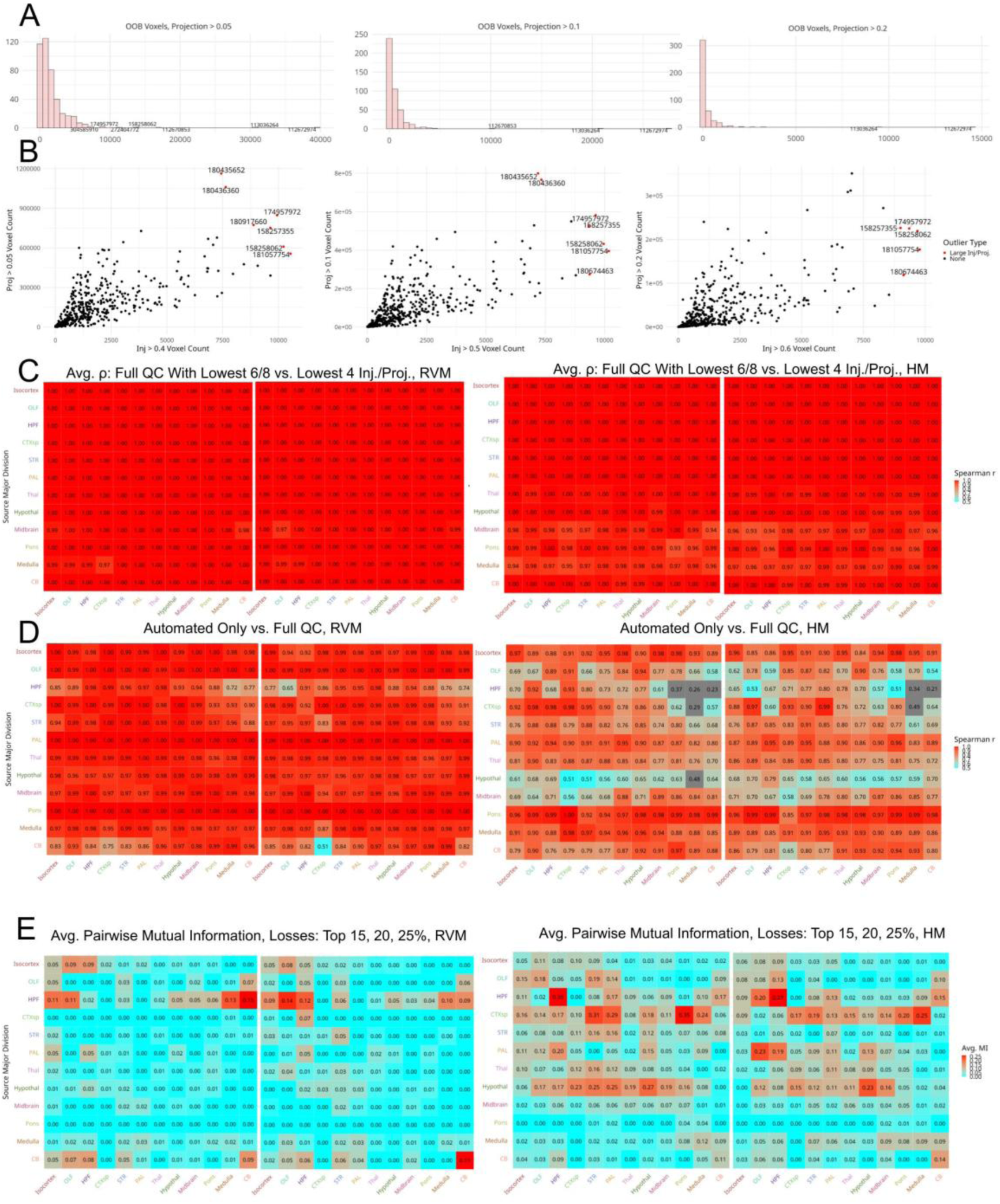
Sensitivity analysis of (A) the out-of-brain outliers based on projection threshold (B) the outliers for injection and projection voxels, based on injection and projection thresholds (C) average pairwise correlation in reconstructed connection strengths within each major division, varying the number of automated lower outliers for injection/projection counts, but keeping all other excluded experiments the same (D) reconstructed connection strengths, when only excluding automated QC failures versus all manual+automated failures (E) average pairwise mutual information in the binarized “losses” in connectivity after QC, when examining the top 15, 20, and 25% of connections.

## Supplementary Information

**Supplementary Methods Figure 1:**
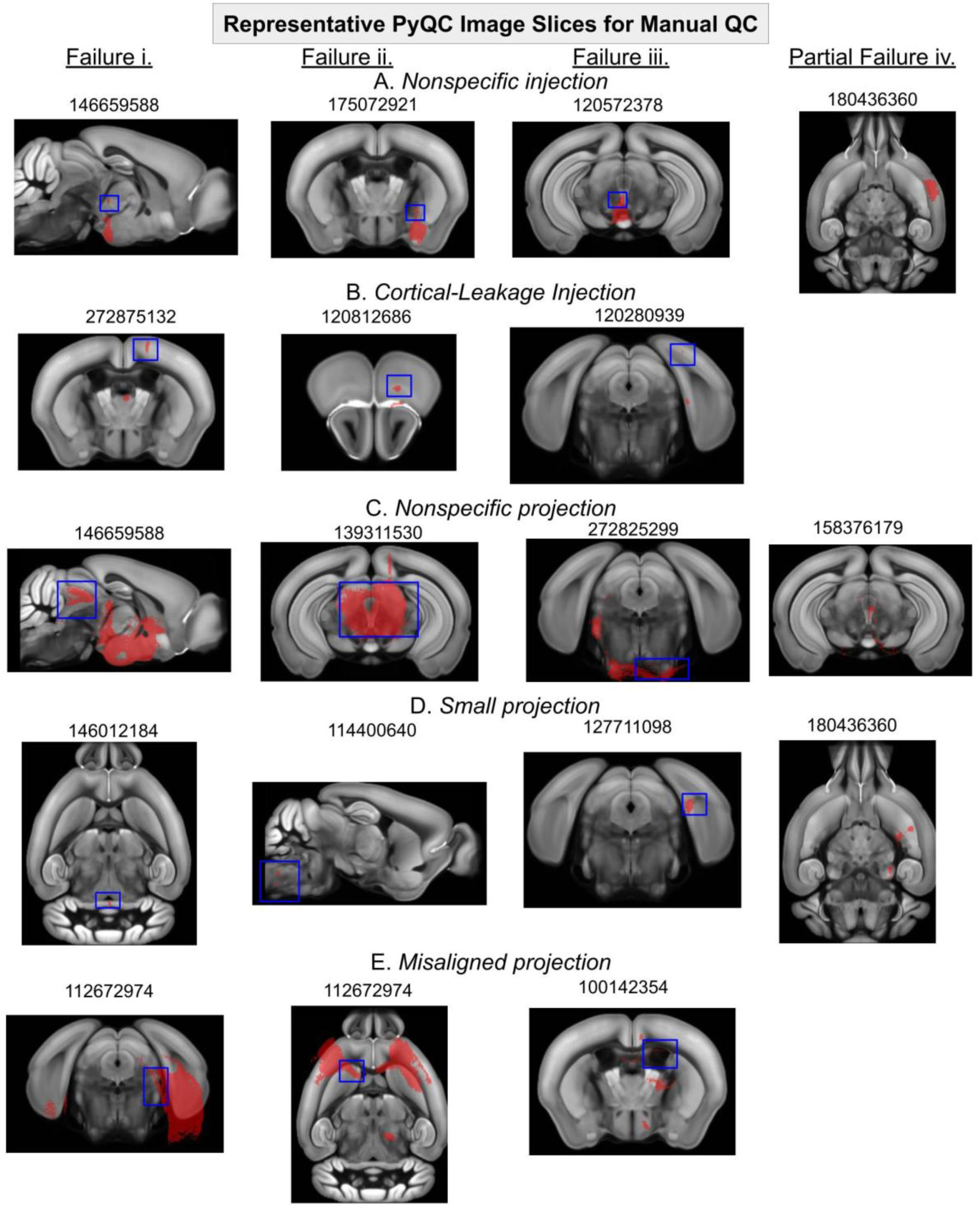
We show representative slice images used in PyQC for our five manual QC modalities, and highlight false-positive or false-negative connectivity patterns in blue. In (A), we show nonspecific injection volumes. We see this failure in (i) the midbrain, following a hypothalamic injection (ii) off-target striatal injection into amygdalar-like areas (iii) nonspecific midbrain injection that infects many regions, including the highly-connected periaqueductal gray (iv) a “passed” experiment that shows a large injection, but is contained within a single cortical region. In (B), we show injection volumes with off-target “leakage” that incorrectly assign cortical connections to the (i) thalamus (ii) olfactory bulb (iii) hippocampus. In (C), we show projection volumes with “nonspecific” connectivity, which could indicate diffuse tracer spreading and “noisy” false-positive connectivity patterns (i) from the hypothalamus to the midbrain and hindbrain (ii) across a variety of midbrain regions (iii) into a number of out-of-brain voxels spanning the edge of the pons (iv) a “passed” midbrain experiment with some evidence of “noisy” spreading to the brainstem. In (D), we show projection volumes that do not spread beyond the injected region, indicating false-negative connectivity patterns to the (i) cerebellum (ii) midbrain (iii) hippocampus (iv) a “passed” striatal experiment that shows some connectivity to other regions, but is not as widespread as we would expect from the striatum. Finally, in (E), we show examples of misaligned projections, which result in the false-positive assignment of connectivity (i) from the hippocampus to the hindbrain (ii) from the corpus callosum to the striatum (iii) within the ventricles.

### Extended Description: Manual QC Failures

We observe a wide variety of experiments removed across major brain divisions and failure modalities (Figure 3C), with the greatest number of removed experiments with injections in the hypothalamus, hippocampus, and midbrain. We see that manual QC is most effective at identifying cortical leaking injections (12 manual only out of 13 total removals), nonspecific injections (12/15), and small projections (12/15) from the injected region (Supplementary Figure 1). When examining the first failure mode for each experiment, we also noticed consistencies across major brain divisions. For example, all n=3 failed cerebellar experiments had small projections. Additionally, for the n=18 failed hypothalamic and midbrain experiments, all but two of the experiments had off-target injections. Nonetheless, for the hippocampal formation, we also observed a mix of connectivity failures, including out-of-brain projections (n=1), cortical leaking injections (n=3), small projections (n=5), and misaligned projections (n=1).

To examine the locations of the lost connectivity, we calculated the number of removed injections (thresholded and binarized) per Allen brain region and the number of removed projections per voxel (Supplementary Figure 2). We then divided this number by the initial counts of injections within each brain region and projections in each voxel to get the ratio of injection experiments removed per region and projection experiments removed per voxel (Figure 3B and 3C). We chose to examine injection removals per region instead of per voxel due to the relative sparsity of injection volumes compared to projection volumes, and due to the importance of injections per region in the HM from Oh et al. (2014). We find that we lose all injections into small cerebellar subregions and brainstem nuclei, which could potentially result in false-negative connectivity to these regions; however, since all other regions have at least one remaining injection, our procedure also has the potential to reduce false-positive connections (Fig. 3B) As expected, we lose 100% of projection voxels outside of the brain and within the medial ventricle, but also lose large proportions (∼75%) of projections to hypothalamic areas (Fig. 3C). Still, we see a loss in projection voxels across the brain, indicating that our QC removals are not specific to a single brain region.

Notably, among our manually excluded experiments, we observed that automated QC identified a large number of experiments with “manual nonspecific” (n=4/7) and “manual misaligned” (n=3/4) projection failures (Supplementary Figure 1). Additionally, of the n=11 experiments with both manual and automated QC failures, n=3 of these experiments failed at least three independent QC criteria (experiment IDs 158257355, 113036264, and 174957972), providing compelling evidence for removal of these experiments and the validity of our criteria. The most extreme example of this was thalamic injection experiment number 174957972, which was automatically flagged for large injection and large projection, as well as manually flagged for large injection and projection volumes.

### Rich Club Algorithm

1. Use *rich_club_bd.m* to calculate the rich club coefficient at each degree.
2. Generate 1000 rewired randomized networks using randmio_dir.m; calculate the rich club coefficient at each degree.
3. For each degree, see the proportion of networks from (2) that have a higher rich club coefficient than the coefficients from (1); use this to define which degrees have a significant number of nodes in a rich club
4. Identify a coefficient defining the “topological rich club” (i.e. whether a node is “rich” or not). This is equal to the mean plus one standard deviation of the degrees with significant rich clubs.

a. Take all the nodes with degrees above this to be “rich club” nodes.
5. For each node with degree k_n_, calculate the percentage of significant degrees that k_n_ is greater than or equal to. This is the rich club percentage, to be visualized in Fig. 5. You can now merge contralateral/ipsilateral nodes, since they will have the same rich club percentage.

### Community Detection Algorithm

1. For each resolution parameter “gamma” from 0.3 to 3, in increments of 0.05:

a. Run the Louvain community detection algorithm (community_louvain.m) 100 times.
b. Find the node x node agreement matrix between the 100 repetitions (using consensus_und.m).
c. Calculate the consensus community assignments from the agreement matrix by using threshold *tau*. The threshold *tau* used to binarize/cluster the agreement matrix is the mean agreement of the ***permuted*** community assignments from (a) (see Lancichinetti & Fortunato, 2012).
2. Write out the community assignments for each gamma.
3. Calculate a gamma x gamma mutual information matrix between the clusterings.
4. Determine which gamma yields the most stable clustering.

a. For mutual information thresholds between 0.9-1 in increments of 0.005:

i. Find the consensus clustering of the gamma x gamma matrix at the given threshold.
b. Calculate the gamma x gamma agreement matrix across all mutual information thresholds.
c. Visualize all values of this matrix above 0.6; heuristically select gamma in the center of the largest “cluster” of agreement matrix values. We chose gamma=1.3, as this was in the middle of a large “cluster” of agreement matrix values in the old and new regionalized voxel connectomes.
5. Run the Louvain clustering with this gamma 100 times; take the consensus clustering as the community assignments for each node.
6. Manually inspect community assignments and ensure that the numbers assigned for each community are consistent across the old versus the new connectomes. If there are two communities that have the same sets of nodes contralaterally vs. ipsilaterally, they can be merged into a single community.

### Sensitivity Analysis of QC Across Parameters

Our QC, and in particular our automated QC, depends on numerous thresholds and cutoffs. To verify the robustness of our approach, we repeated our QC across a number of different parameter choices. First, for our automated QC, we varied the binarization thresholds used for the injection and projection data (Supplementary Figure 10A). We see differences in the number of experiments identified as outliers for out-of-brain voxels based on the projection threshold, ranging from n=7 to n=2 removed experiments, but note that the top two experiments for out-of-brain voxels are robustly identified across all thresholds. We also see a minimal difference in the number of upper outliers for overall injection or projection voxel counts across injection (0.4, 0.5, 0.6) and projection (0.05, 0.1, 0.2) thresholds, with n=5 out of the n=7 excluded experiments remaining the same across thresholds (Supplementary Figure 10B). We also note that reconstructed connection strengths after full QC are robust to differences in the number of automated lower outliers (6 and 8 per inj. or proj., versus the original 4), as confirmed by high average Spearman correlations of connection strengths in each major division in the RVM and HM (min ρ=0.94 in HM; 0.97 in RVM) (Supplementary Figure 10C).

Next, we visualized the correlation in reconstructed connectivity strengths after only removing experiments with automated QC versus all experiments (Supplementary Figure 10D). Across both the RVM (min ρ=0.65) and the HM (min ρ=0.21), the hippocampus shows the biggest difference in connectivity between the automated-only versus the full removals, indicating the importance of our manual QC in detecting the hippocampus-medulla and hippocampus-cerebellum losses in connectivity. Additionally, within the RVM, most other regions show limited differences in automated vs. full QC, whereas the HM is far more sensitive to manual QC, particularly in the hypothalamus.

Finally, we conducted a sensitivity analysis on the binarization threshold to see which binary connectivity “losses” post-QC versus pre-QC were the most consistent within the top 15, 20, and 25% of connections. (Supplementary Figure 10E). We find in the RVM that the connectivity losses with the highest average mutual information are hippocampus-medulla and hippocampus-cerebellum, indicating that these losses in connectivity are stable across binarization thresholds. However, for the HM, although the mutual information of these corresponding changes is well above zero, we see the most consistent connectivity losses in hippocampus-hippocampus connection and cortical subplate to pons connections, highlighting differences in how these models respond to QC.”

We also note that manual QC depends on the binary threshold used to generate the QC images. To verify that our manual QC failures were threshold-independent, V.N. examined all injection and projection images at a lower binarization threshold of 0.01, and did not observe notable differences in the visually-identified experimental “failures.” We include these images within the Supplementary Data.

